# A pangenomic atlas reveals that eco-evolutionary dynamics shape plant pathogen type VI secretion systems

**DOI:** 10.1101/2023.09.05.556054

**Authors:** Nathalie Aoun, Stratton J. Georgoulis, Jason K. Avalos, Kimberly J. Grulla, Kasey Miqueo, Cloe Tom, Tiffany M. Lowe-Power

## Abstract

Soil-borne *Ralstonia solanacearum* species complex (RSSC) bacteria disrupt rhizosphere and endophytic microbial communities as they invade roots and fatally wilt plants. RSSC pathogens secrete antimicrobial toxins using a type VI secretion system (T6SS). To investigate how evolution and ecology have shaped pathogen T6SS biology, we analyzed the T6SS gene content and architecture across the RSSC pangenome and their evolutionarily relatives. Our analysis reveals that two ecologically similar Burkholderiaceae taxa, xylem pathogenic RSSC bacteria and *Acidovorax*, have convergently evolved to wield large arsenals of T6SS toxins. To understand the mechanisms underlying genomic enrichment of T6SS toxins, we compiled an atlas of 1,069 auxiliary (“*aux*”) T6SS toxin clusters across 99 high-quality RSSC genomes. We classified 25 types of *aux* clusters with toxins that predominantly target lipids, nucleic acids, or unknown cellular substrates. The *aux* clusters were in diverse genetic neighborhoods and had complex phylogenetic distributions, suggesting frequent horizontal gene flow. Phages and other mobile genetic elements account for most of the *aux* cluster acquisition on the chromosome but very little on the megaplasmid. Nevertheless, RSSC genomes were more enriched in *aux* clusters on the megaplasmid. Secondary replicons like megaplasmids often evolve more rapidly than the more evolutionarily stable chromosome. Although the single ancestral T6SS was broadly conserved in the RSSC, the T6SS was convergently lost in atypical lineages with vectored transmission. Overall, our data suggest dynamic interplay between the lifestyle of soil-transmitted RSSC lineages and the evolution of T6SSs with robust arsenals of toxins. This pangenomic atlas poises the RSSC as an emerging, tractable model to understand the role of the T6SS in shaping pathogen populations.

## Introduction

To defend their habitat or colonize new niches, bacteria often attack competing microbes by secreting toxic molecules and proteins (1, 2). Many gram-negative bacteria use a molecular weapon known as the type 6 secretion system (T6SS) to deliver toxic protein effectors to target cells (3, 4). T6SS effectors bind to the tip of a multimeric, spear-like projectile (5–7). Upon contracting, the sheath forcibly ejects the projectile, puncturing nearby cells and delivering the associated effectors directly to the intracellular space of target cells within range (3, 8). T6SS effectors kill target cells by degrading cellular components including DNA, lipid membranes, and bacterial cell wall polymers. To prevent damage to kin cells, each T6SS effector has a cognate immunity protein that prevents toxicity, usually by directly binding the effector (8, 9).

Most T6SSs include a double membrane spanning complex (TssJLM) which recruits a baseplate complex (TssEFGK) and a spike complex (VgrG heterotrimer, PAAR sharp tip, effectors) (10–13). Hcp then polymerizes to form the shaft of the projectile, and repeating units of TssBC form a sheath around the shaft that contacts to fire the projectile complex. After firing, TssH disassembles the TssBC sheath, allowing the monomers to reassemble again in a new complex (14, 15). While just one copy of most *tss* genes is required for a functional T6SS, T6SS^+^ genomes usually encode multiple copies of *vgrG*, typically collocated with the cognate effector gene, immunity gene, and genes encoding the PAAR sharpening tip protein and any adapter proteins (13, 16). More copies of *vgrG* in a genome implies a greater diversity of effectors, potentially allowing the T6SS to be deadly against a greater diversity of targets. Clusters of *vgrG* and associated genes may be located with the other *tss* genes or in other locations termed “T6SS auxiliary clusters” (“*aux* clusters”) (17, 18). Horizontal gene transfer has previously been implicated in the spread of *aux* clusters between bacterial genomes (19–22), contributing to convoluted evolutionary patterns of gain and loss that result in diverse T6SS effector arsenals. Even closely related strains often have non-identical T6SS effector repertoires, allowing interstrain competition within a bacterial species (23, 24). T6SS-mediated interstrain competition has been shown to contribute to the outcome of host colonization when two or more strains of the same species attempt to colonize a new host (25). Large-scale analyses suggest that plant-associated and human-associated bacteria are more likely to have a T6SS (26).

Plant pathogens in the *Ralstonia solanacearum* species complex (RSSC) are known to have a functional T6SS (27–29). RSSC pathogens cause severe wilt disease on diverse plants, including economically important crops like potatoes, tomatoes, and bananas. Like many host-associated bacteria, RSSC plant pathogens must successfully transition between disparate ecosystems to complete their disease cycle. RSSC bacteria persist in soil or water and chemotax through the microbially dense rhizosphere to infect a new host (30). *Ralstonia* enter the roots and spread throughout the xylem, growing to such a high density that the mass of cells and biofilm physically block water transport, wilting the plant (31, 32). Transcriptomic analyses indicate that the T6SS is upregulated when RSSC pathogens reach quorum, as in an infected plant (33). T6SS-inactive *Ralstonia* mutants have pleiotropic phenotypes, including altered biofilm and motility, which has made it difficult to define the precise ecological role for the RSSC T6SS.

We hypothesized that intermicrobial competition during plant colonization has shaped the evolution of RSSC pathogens and their T6SS weapons. Here, we used an evolutionary genomics approach to shed light on the eco-evolutionary dynamics between the lifestyle of RSSC strains and their T6SS arsenals. We carried out a pangenome analysis of T6SS-related genes on diverse genomes in the RSSC compared to Burkholderiaceae family relatives. We found that that plant pathogenic *Ralstonia* genomes are evolutionarily enriched in T6SS effectors. Scrutinizing the diversity and distribution of T6SS effectors in the RSSC, we found a complex distribution of T6SS *aux* clusters suggesting that *aux* clusters are frequently gained and lost. More of the identified T6SS gene clusters were present on the megaplasmid than the main chromosome. Intriguingly, most of the chromosomally located *aux* clusters were associated with mobile genetic elements, while there was rarely evidence of mobile genetic elements associated with megaplasmid-located *aux* clusters. We infer that the T6SS is an ancestral trait in the RSSC whose genetic architecture is shaped by complex ecological and evolutionary processes. Our analyses reveal that the T6SS is a dynamic weapon intimately linked to the evolutionary success of soilborne *Ralstonia*, a plant pathogen of global concern.

## Results

### Bacteria in the family Burkholderiaceae vary in their T6SS gene content

We used an evolutionary framework to explore T6SS gene content among the RSSC relative to their evolutionary neighbors in the Burkholderiaceae family. Using the genome taxonomy database (GTDB) (34–39), we identified complete genomes of 288 Burkholderiaceae species (Table S1, S2). We used BLASTp searches to estimate the abundance of T6SS genes in the Burkholderiaceae genomes and a curated set of 99 high-quality RSSC genomes. To estimate the number of secretion systems per genome, we averaged the number of BLASTp hits for the core T6SS components: TssABCEFGHJKLM and Hcp (40). The majority of the Burkholderiaceae species representatives encoded a T6SS (62.4%) (Figure S1, Table S1). The three species representatives of the RSSC wilt pathogens (*R. solanacearum* IBSBF1503, *R. pseudosolanacearum* GMI1000, and *R. syzygii* PSI07), each encoded a single copy of the core T6SS genes. In contrast, there were multiple T6SSs encoded in approximately 27% of the Burkholderiaceae genomes; between three to six sets of T6SS core genes were identified in genomes in the *Burkholderia*, *Trinickia*, *Caballeronia*, *Variovorax*, *Pseudoduganella*, *Massilia*, and *Paraburkholderia* genera (Figure S2; Table S1), which is consistent with previous reports for some of these taxa (41–45).

### Genomes of RSSC plant pathogens are enriched in *vgrG* genes

T6SS effector/immunity pairs are often encoded in gene clusters with their corresponding *vgrG,* any adaptors that mediate effector translocation, and any genes encoding the PAAR sharpening tip protein (46–48). Because VgrG proteins have conserved sequences, we estimated the abundance of effector/immunity pairs by searching genomes for *vgrG* copies. We carried out multiple low-stringency BLASTp searches for *vgrG* alleles in the 288 Burkholderiaceae species representatives and the 99 high-quality RSSC genomes. As expected, most of the T6SS^null^ genomes lacked intact *vgrG* genes (Figure S2). Generally, there was a positive correlation between the copy number of T6SS core genes and *vgrG* alleles (Figure 1A, S2).

**Figure 1.**
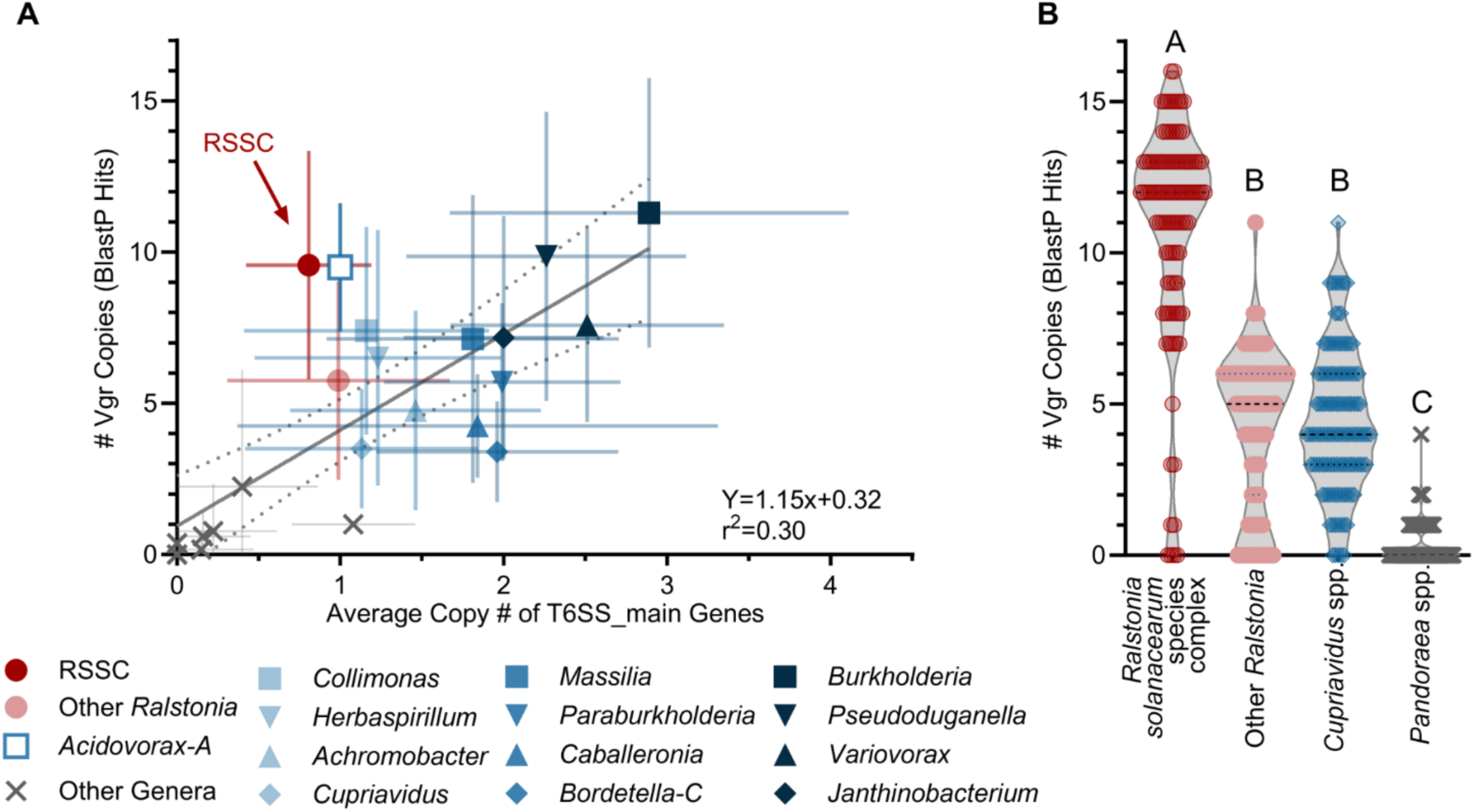
RSSC genomes are enriched in *vgrG*-linked effector/immunity clusters. (**A**) The number of *vgrG* alleles and T6SS core components (TssABCEFGHJKLM and Hcp) were compared across species within the Burkholderiaceae family. Using a custom genome database in KBase, we identified genes with BLASTp searches. The Grey “X” symbols indicate the nine genera with few-to-no T6SS genes (*Acidovorax, Alcaligenes, Bordetella, Comamonas, Hydrogenophaga, Pandoraea, Polaromonas, Polynucleobacter, Rhodoferax*). Error bars indicate SD, and dashed lines indicate the 95% confidence bands around the linear regression. (**B**) The number of *vgrG* genes were estimated in RSSC genomes (n=99) relative to closely related species/genera: other *Ralstonia* spp. (n=70), *Cupriavidus* spp. (n=120), and *Pandoraea* spp. (n=75). *vgrG* alleles were identified via BLASTp on KBase. Letters indicate p<0.0001 by Kruskal-Wallis multiple comparisons test.

Two taxa with single copies of the core T6SS genes were enriched in *vgrG* genes: the plant-pathogenic RSSC and the genomo-genus Acidovorax_A (Figure 1A). Acidovorax_A is a genomo-genus defined by GTDB (34–39) that contains two xylem-infecting plant pathogen species, *Acidovorax avenae* and *Acidovorax citrulli*. To investigate whether the RSSC had an evolutionary enrichment of *vgrG* copies, we compared the number of *vgrG* genes in the RSSC to other species in the *Ralstonia* genus (n=70) and the closely related genera *Cupriavidus* (n=120) and *Pandoraea* (n=75). Plant-pathogenic RSSC genomes typically had over twice as many *vgrG* copies as their close relatives (median of n=12 per genome; Figure 1B). Most *Pandoraea* spp. did not encode *vgrG*. *Cupriavidus* and the non-RSSC *Ralstonia* encoded a similar number of *vgrG* copies (medians of n=4 or n=5 per genome, respectively) (Figure 1B).

### A single T6SS subtype is largely conserved among plant pathogenic *RSSC*

We used synteny analyses to investigate the organization of T6SS genes among the plant pathogenic RSSC. The genomes we investigated have the same T6SS subtype, T6SS^i4b^ (49, 50) (Figure 2A), encoded on the 2.1 Mb secondary replicon known as the megaplasmid (51). The RSSC T6SS has two conserved regions interspersed between three variable regions that mainly contain *vgrG*-linked effector/immunity clusters and small transposable elements (Figure 2A-B).

**Figure 2.**
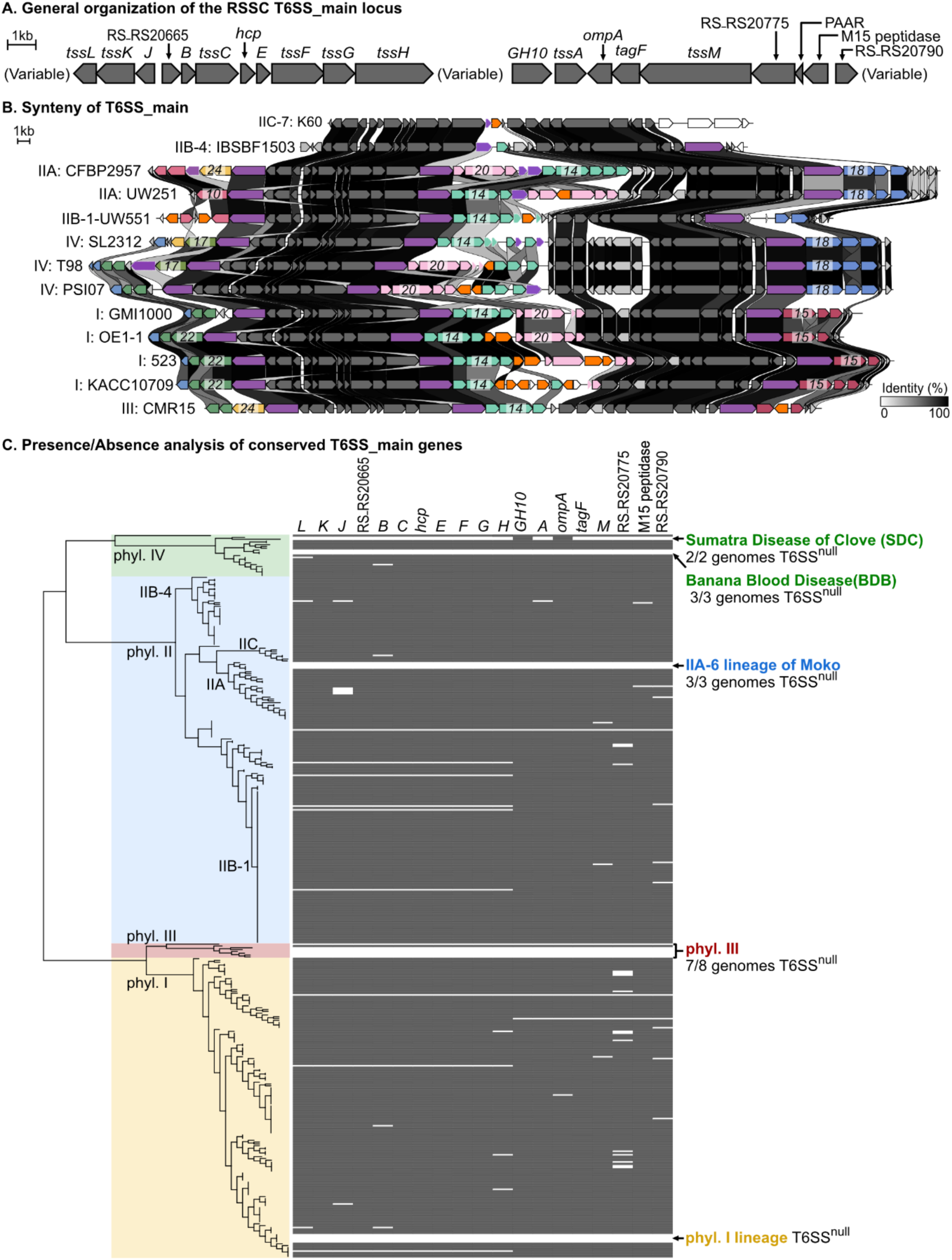
Although the T6SS is broadly conserved, multiple RSSC lineages lack the T6SS. Most RSSC encode a type 6 secretion system (T6SS) with a conserved gene order with three variable regions and two conserved regions. (**A**) shows the conserved genes and (B) displays the synteny of the locus across phylogenetically diverse RSSC strains. Core T6SS genes encoding structural components and associated genes are shown as grey arrows, VgrG spike proteins are shown as purple arrows, and IS elements and transposases as orange arrows. Numbers superimposed over genes in the variable regions identify which auxiliary effector/immunity clusters are encoded at each spot. Linkages are drawn between homologous genes. Synteny and global amino acid identity was visualized with Clinker and aesthetics were adjusted in Affinity Designer. (**C**) Left: A phylogenetic tree of 398 RSSC genomes was created with the KBase “Species Tree” app, which compares 49 conserved bacterial genes. Right: The presence/absence of core T6SS genes based on BLASTp analysis. Grey rectangles indicate at least one homolog was identified using the KBase BLASTp v2.13.0+ app with percent identity cut-offs of > 15% and aligned-length cut-offs of > 20%. The data was visualized on iTOL. Lineages that lack the T6SS are indicated.

Excluding the three *vgrG* paralogs, we identified 19 core genes in the T6SS locus (Figure 2A). We BLAST-searched the 19 genes against the 288 Burkholderiaceae species representatives. Although the structural genes were well-conserved, we observed that six of the core T6SS^i4b^ genes were variably present in 21.4-70.9% of the other T6SS-containing genomes: RS_RS20665, *GH10* (glycosyl hydrolase 10), *ompA*, *tagF*, RS_RS20775, and the M15 peptidase (Figure S1). One gene encoding a hypothetical protein, RS_RS20790, was exclusively found in *Ralstonia* genomes (Figure S1).

### Several RSSC lineages have lost T6SS

Upon searching 398 publicly available RSSC genomes, we identified several lineages that lacked the T6SS (Figure 2C). The T6SS^null^ lineages include the two insect and mechanically vectored phylotype IV lineages that cause Sumatra Disease of Clove (SDC) and Blood Disease of Banana (BDB), one of the lineages that causes Moko Disease of Banana (phylotype IIA-6), one phylotype I lineage, and all but one of the eight phylotype III strains with sequenced genomes. Of these T6SS^null^ lineages, the SDC lineage strains encode the *gh10* and *ompA* genes, suggesting that the other T6SS genes were lost. We also identified multiple genomes that lacked the conserved *tssL*-to-*tssH* and the *GH10*-to-RS_RS20790 region, (n=30 and n=24, respectively; Figure 2C, Table S2). There were sporadic genomes that lacked a BLASTp hit for certain core genes, but these could be false negatives based on carrying out BLASTp based searches on draft genomes of varying quality.

To infer whether the T6SS^i4b^ had been lost in the T6SS^null^ lineages, we carried out BLAST searches and synteny analyses with all the T6SS gene clusters identified in RSSC and the non-RSSC *Ralstonia*. Of the T6SS^+^ non-RSSC *Ralstonia*, 98% of the genomes encoded a T6SS^i4b^ with the same gene organization except for the absence of the GH10-encoding gene (Figure S3). The simplest explanation for the phylogenetic pattern is that the T6SS^i4b^ is ancestral to the genus *Ralstonia* and has independently been lost in multiple lineages of the RSSC and non-RSSC *Ralstonia*.

### The RSSC pangenome encodes dozens of auxiliary T6SS effector/immunity clusters

Because RSSC genomes contain a median of 12 *vgrG* genes and only three *vgrG* paralogs are encoded in the main T6SS^i4b^ locus, we hypothesized that there were additional T6SS loci on the chromosome or megaplasmid. We performed an iterative synteny analysis of the gene neighborhoods of all *vgrG* alleles in the 99 high-quality RSSC genomes, which allowed us to infer the presence of 1069 auxiliary effector/immunity (*aux*) clusters that we classified into 25 types: *aux1-aux25* (Table S3). Furthermore, we used synteny analyses to map the *aux* clusters to their locations on the chromosome, megaplasmid, or small accessory plasmids. Figures S4-S24 show an atlas for each *aux* cluster, depicting the phylogenetic distribution, gene organization and annotations, and genomic locations.

The *aux* clusters contain between three and 14 genes and vary in size from 2.27 kb (*aux17*) to 16.6 kb (*aux16*). We used the NCBI Conserved Domain Database and PaperBlast to infer the function of each *aux* cluster gene (Figures S4-S24). Although there is a conserved standalone PAAR domain gene in the T6SS main, small PAAR domain-containing genes were also encoded in some or all structural variants of five *aux* types (*aux1, aux2, aux5, aux7, aux9*). While some T6SS effectors bind directly to VgrG spike proteins, others must form complexes with adaptor proteins (52). We identified adaptors with DUF1795 (24) and DUF2169 (53) in three *aux* types and DUF4123 domains (54) in four *aux* types. Several of the *aux* clusters include genes containing polymorphic effectors with recombination hotspot (RHS) domains (*aux3, aux4,* and *aux12*), DUF4150 PAAR-like superfamily domains (*aux8*, *aux13*, and *aux16*) (55), marker-for- type-six (MIX_III) domains (*aux14* and *aux20*) (56), found-in-type-six (FIX) domains (*aux10*, *aux17*, *aux22*, *aux23*, and *aux24*) (17), and FIX-like domains (*aux19* and *aux21*). Finally, genes also included other domains that have been previously identified in T6SS effectors (DUF3274, DUF2235, DUF2875, and DUF6531) (57–59) and immunity proteins (DUF1910, DUF1911, DUF3304, and Sel1 repeats) (17, 18, 57) (Figures S4-S24). We identified putative arrays of immunity genes or orphan immunity genes in structural variants in 13 of the *aux* types: *aux2*, *aux3*, *aux4*, *aux5*, *aux6*, *aux8*, *aux11*, *aux12*, *aux14*, *aux15*, *aux18*, *aux22*, and *aux23*.

The largest two clusters, *aux8* and *aux16*, are atypical because they were the only ones that contain genes upstream of the *vgrG* gene: one DUF4124-encoding gene and 1-2 ankyrin repeat-encoding genes (Figures S11 and S19). All genes in *aux1*-*aux24* were arranged unidirectionally, but *aux25* had an atypical three-gene layout of *vgrG*, a 2.3 kb hypothetical gene, and then a conserved inverted gene encoding a PvdO family nonheme iron enzyme (Figure S24).

### RSSC-encoded T6SS effectors target diverse substrates, including lipids and DNA

T6SS effectors damage important cellular components in target cells. We used the NCBI Conserved Domain Database (60) and PaperBlast (61) to infer the mode-of-action of the effectors (Figure 3, Table S3). The effectors from nine *aux* clusters were lipases of previously defined families (62): Tle1 (*aux6* and *aux11*), Tle3 (*aux2, aux7,* and *aux9*), Tle4 (*aux1* and *aux15*), Tle5 (*aux6*), and a lipase with a novel domain architecture (*aux18*). The effectors of seven *aux* clusters were nucleases with HNH nuclease domains (*aux3*, *aux12*, and *aux16*), GHH2 nuclease domains (*aux8*), or PoNe nuclease domains (*aux10*, *aux19*, and *aux21*). The polymorphic RHS effectors in *aux4* varied in the C-terminal effector, containing either ATP-targeting (p)ppApp synthase or actin-ADP ribosylase. For the remaining nine clusters, we could infer which gene was the effector based on the presence of known polymorphic effector domains for eight of the clusters (*aux3, aux14, aux17, aux20, aux22, aux23, aux24,* and *aux25),* but we could not identify domains or motifs that hint at the mode-of-action (Figure 3E). For the final cluster, *aux13*, the effector might be either the MAEBL-domain protein or one of the three hypothetical proteins.

**Figure 3.**
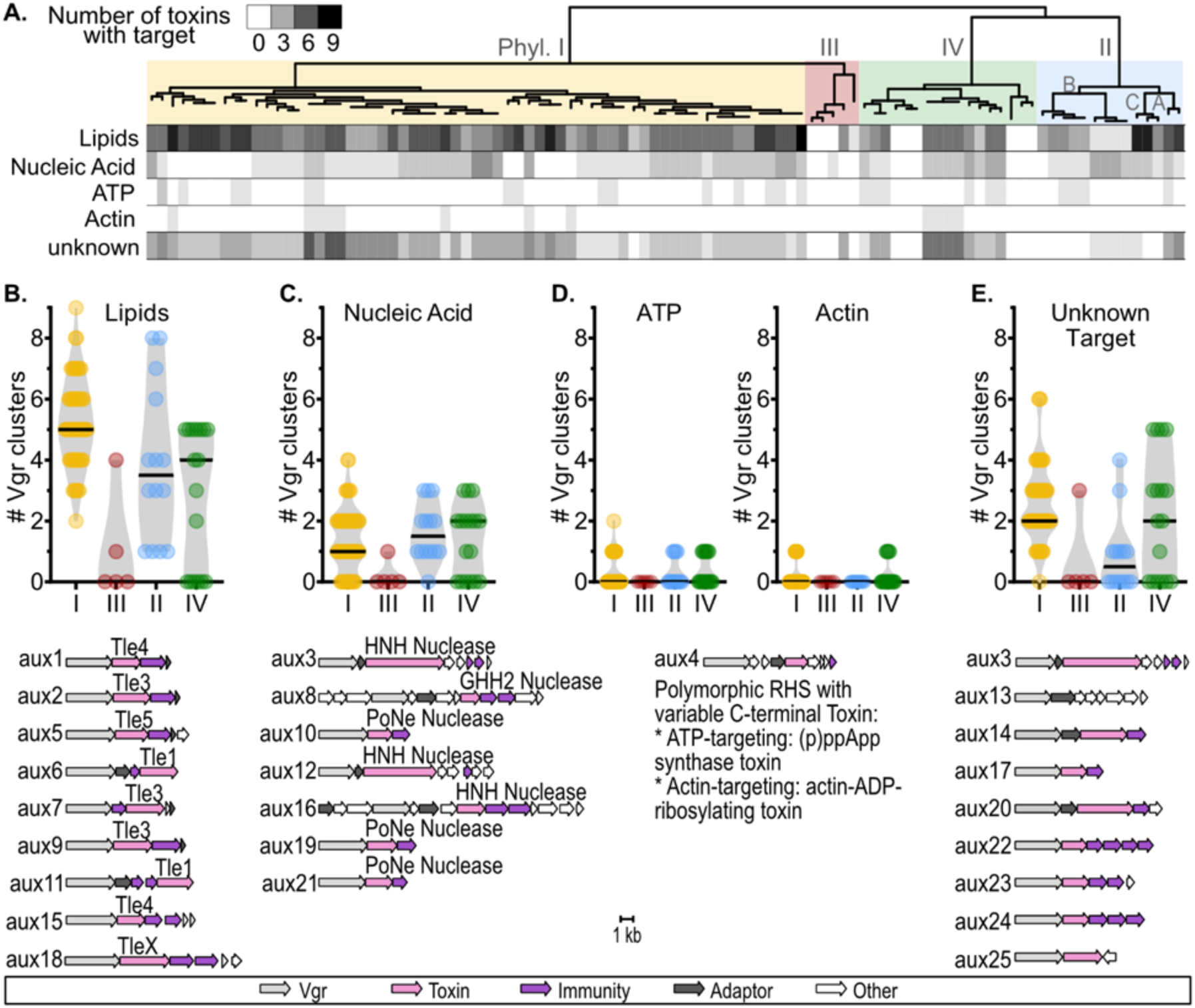
RSSC T6SS effectors are predicted to target lipids, nucleic acids, ATP, actin, and unknown targets. To identify effectors and analyze their sequence for enzymatic domains, all Vgr-linked sequences were queried against NCBI Conserved Domain Database and PaperBlast. (A) Phylogenetic analysis of the effector profiles. (B-E) Comparison of clusters with (B) unknown effectors, (C) Lipases, (D) Nucleases, and (E) *aux4*, a polymorphic effector cluster that varies in the effector target.

We investigated whether RSSC lineages varied in the abundance of lipase, nuclease, ATP/Actin manipulating, and unknown-targeting effectors (Figure 3). All T6SS^+^ RSSC genomes encoded at least one lipase. Phylotype I and IIC lineages encoded the most lipases (median of 5 and 8, respectively) (Figure 3A- B). Phylotype IIB-1, which contains the clonal pandemic lineage (regulated as a U.S. Select Agent under the name “*R. solanacearum* R3Bv2”)(63), had the fewest lipases of the T6SS^+^ genomes with only a single lipase per genome. Nucleases were common, but 17% of T6SS^+^ strains lacked any obvious nucleases. Sporadic phylotype I strains, the IIB-1 lineage, and the IV-8 lineage had more nucleases than the other lineages (Figure 3A). Genomes with more nucleases tended to have fewer lipases. The ATP and actin-targeting effectors are rare and sporadically distributed. None of the analyzed phylotype II genomes encoded an actin-targeting effector. Most of the T6SS^+^ genomes (91%) had one or more effectors with unknown targets.

### T6SS gene conservation varies among RSSC clades

The copy number of *aux* types ranged from zero to three copies among the RSSC genomes (Figure S25). Some *aux* types were never found with more than a single copy per genome. *aux2, aux5,* and *aux7* were sometimes found in two or three copies, and *aux1, aux4, aux6, aux9,* and *aux17* were sometimes found in two copies across the RSSC genomes. The *aux10, aux11, aux12, aux18, aux19*, *aux21*, *aux24* and *aux25* were found in only 20% of the RSSC genomes. Meanwhile, *aux1*, *aux*2, *aux*3, *aux*5, *aux*6, *aux*7, *aux*14, and *aux*15 were found in more than 50% of the RSSC genomes (Figure S25).

To compare the phylogenetic distribution of different *aux* types, we visualized the copy number of each *aux* type across an RSSC phylogenetic tree (Figure 4C and Figure S26). To organize the *aux* types based on their patterns of presence in each genome, we calculated the Bray-Curtis dissimilarity metric based on the *aux* cluster content of each strain and used the *aux* cluster content dissimilarity matrix to draw a dendrogram (Figure 4C, Table S4). The dendrogram divides *aux* clusters into two groups: one group where *aux* clusters are highly prevalent across the strains and another group where *aux* clusters are less common (Figure 4C). *aux1-3*, *aux6*, and *aux14* were present in at least one genome of every phylotype, excluding the mostly T6SS^null^ phylotype III. *aux4, aux9*, *aux10*, and *aux22* were found in each of the remaining three mostly T6SS^+^ phylotypes. In contrast, some *aux* types were restricted to particular phylotypes: *aux13* and *aux19* were present only in phylotype I, while *aux12* and *aux25* are only found in phylotype II. Additionally, there were species-specific patterns for some *aux* clusters. *aux15* and *aux22* are present in *R. pseudosolanacearum* (phylotype I and III) but absent in *R. solanacearum* (phylotype II) and *R. syzygii* (phylotype IV). *aux24* and *aux25* are rare and were only identified in two and one RSSC genomes, respectively (Figure 4C). As a result of these highly variable patterns of *aux* cluster distribution, closely related strains typically have overlapping but non-identical *aux* cluster repertoires (Figure 4C).

**Fig 4.**
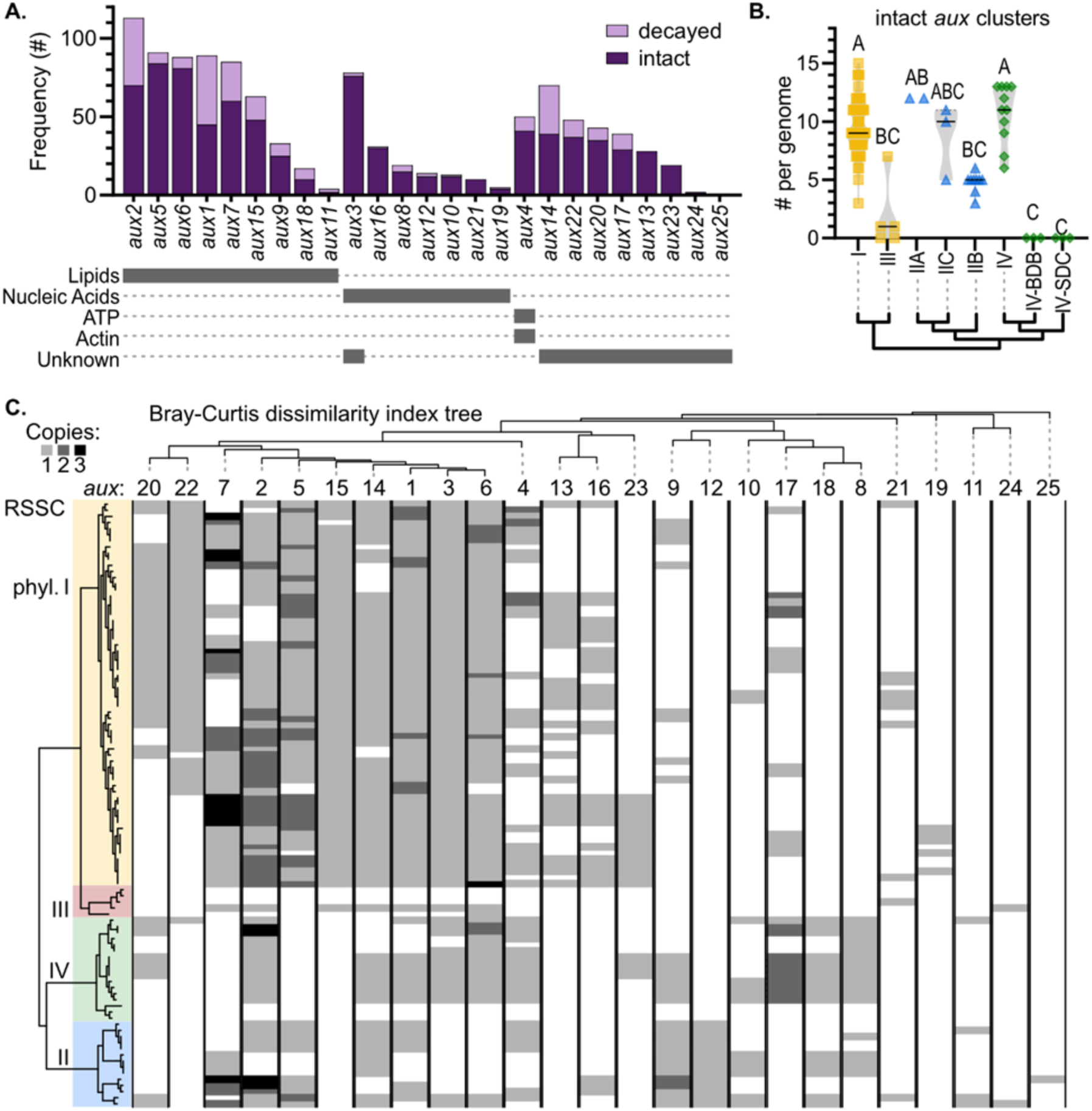
RSSC pathogens vary in their repertoires of T6SS auxiliary effector/immunity clusters (“*aux*”). (**A**) Synteny analysis classified 1069 *vgr*-linked clusters into 25 subtypes (*aux1-aux25*) that varied in their prevalence from common to rare. Clusters were classified as “decayed” (pink) if genes were pseudogenized or were disrupted by transposons and were classified as “intact” (purple) if they lacked obvious mutations. (**B**) Relative abundance of intact *aux* clusters across major phylogenetic divisions of the RSSC. Lines indicate the median, and a cladogram shows the relative evolutionary relationships of the groups. Letters indicate significance groups by Kruskal-Wallis test with multiple comparisons (p<0.05). (**C**) Presence of each cluster (*aux1-aux25*) was determined across complete or nearly complete RSSC genomes by a combination of BLASTp and Clinker analyses. The left tree represents phylogeny of the *Ralstonia* genomes based on analysis of 49 genes. The top tree represents Bray-Curtis dissimilarity of the 25 *aux* clusters across the analyzed genomes. The phylogenetic tree was built with the Kbase SpeciesTree app, Bray-Curtis dissimilarity indices were calculated with the ‘vegan v.2.6-2’ package in RStudio (2023.03.0+386), and the visualization was created with iToL.

### *aux* clusters are enriched on the megaplasmid

In bacterial genomes, secondary replicons like the RSSC megaplasmid often contain more accessory genes which evolve more quickly than chromosomal genes (64–66). Because *aux* clusters are part of the RSSC accessory genome, we hypothesized that they would be more common on the megaplasmid. We found that *aux* clusters were enriched on the megaplasmid, accounting for 71.6% of the 1069 *aux* clusters that we classified (Figure 5, Figure S27, and Table S3). Considering that the megaplasmid is smaller than the chromosome (approximately 2.1 Mb and 3.5 Mb, respectively), *aux* cluster density is dramatically higher on the megaplasmid. Although most *aux* types are more common on the megaplasmid, certain *aux* types were more common on the chromosome: *aux2, aux8, aux21, aux19, aux4, aux17,* and *aux13* (Figure 5B and Table S3).

**Figure 5.**
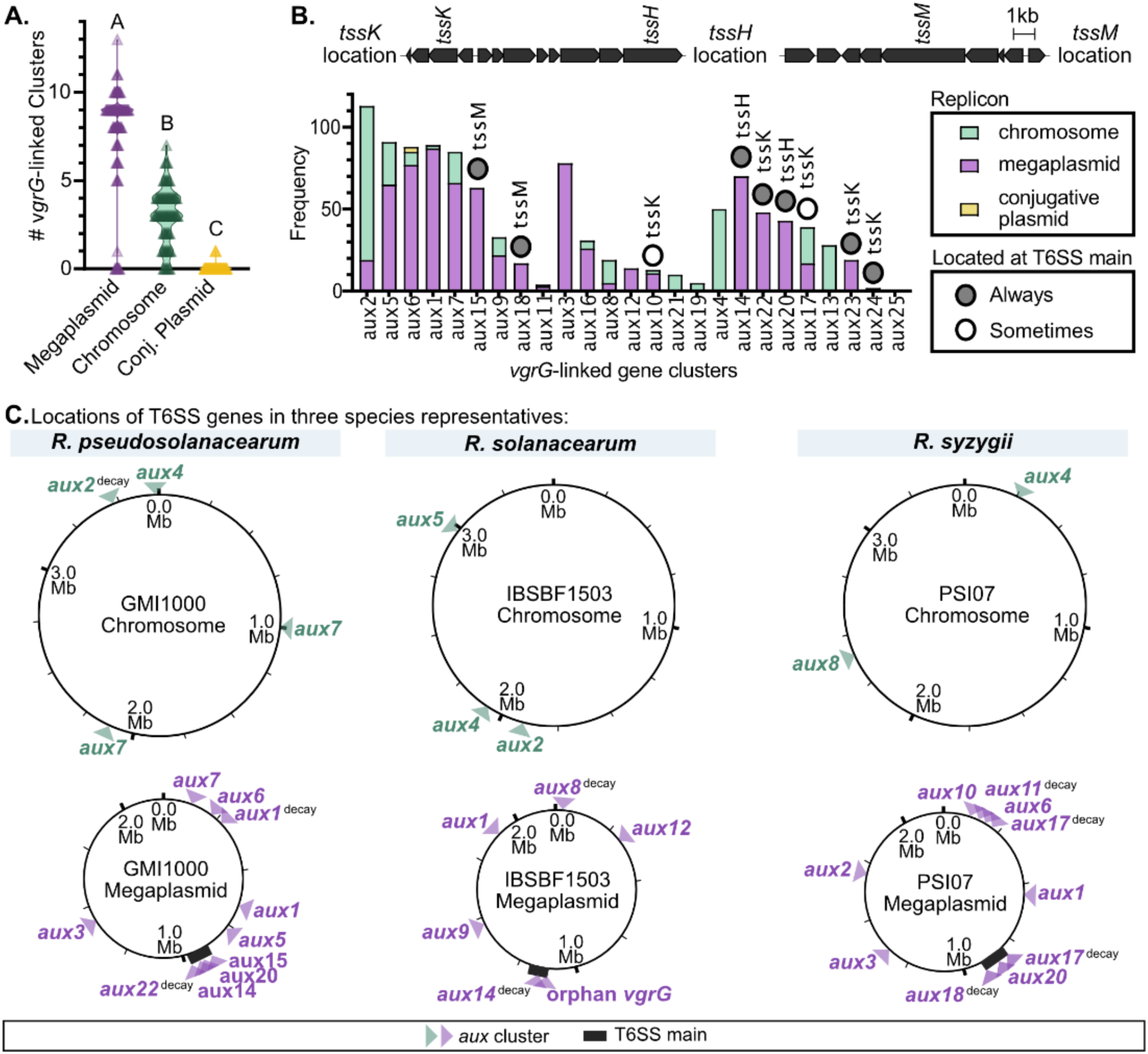
T6SS *aux* clusters are enriched on the megaplasmid. (**A**) The number of *aux* clusters on each replicon (the chromosome, megaplasmid, and accessory conjugative plasmids) per RSSC genome. Letters indicate p<0.0001 by Brown-Forsythe and Welch ANOVA with Dunnett’s T3 multiple comparisons test. (**B**) The occurrence of each *aux* cluster type on the replicons was identified. *aux* clusters are sorted by the substrate that the effector degrades (See Fig 3-4). Nine *aux* clusters were always (dark grey circles) or sometimes (white circles) located in the T6SS main locus. The three locations where *aux* clusters are found in the T6SS main locus are indicated above the graph. (**C**) The locations of *aux* clusters on the chromosome (green arrowheads) and megaplasmid (purple arrowheads) of RSSC species representatives were visualized. The species representatives (*R. pseudosolanacearum* GMI1000, *R. solanacearum* IBSBF1503, and *R. syzygii* PSI07) lack accessory conjugative plasmids.

To determine whether there are genomic islands where *aux* clusters are located, we used Clinker synteny analyses to map the specific location of each of the 1069 *aux* clusters. Except for the T6SS main locus, there were no obvious hotspots where *aux* clusters were located (Figure 5C and panel C of Figure S4-S24). The three variable regions of the T6SS main locus each contained *aux* clusters that had the same superfamily of effectors. The variable region downstream of *tssM* only contained *aux15* or *aux18*, which had lipase effectors (Figure S18). The variable region between *tssH* and *tssA* exclusively contained *aux14* and/or *aux20*, which have a MIX_III domain effector (Figure S17). All *aux* clusters found downstream of *tssJKL* contained effectors with FIX domains, including *aux22-24*, which were exclusively found at T6SS main, and *aux10* and *aux17*, which were also found associated with the T6SS main cluster as well as additional loci (Figure S13, S20). Certain *aux* types were located in consistent genomic loci while others were highly variable. For instance, *aux3* is found in 79% of surveyed RSSC genomes and is always found in the same locus on the megaplasmid (Figure S6). In contrast, a*ux2* is found in 84% of our genome set, but it is located in nine distinct loci (Figure S5).

Some of our RSSC genomes included accessory plasmids that encoded two different aux types: *aux6* and *aux11*. However, both of these *aux* types were also found on the chromosome or megaplasmid in other RSSC genomes (Figure S9, S14).

### Loss-of-function mutations are common among *aux* clusters

To infer the role of gene loss in the evolutionary history of *aux* clusters, we used Clinker synteny analyses to identify apparently functionally “intact” *aux* clusters and “decayed” clusters with one or more putative loss-of-function mutations (Figure S28). Loss-of-function mutations included large deletions, frameshifts, premature stop codons, or disruption by insertion sequence (IS) elements (Table S3). Of the 1069 *aux* clusters classified, about 23.5% contained one or more *vgrG*, effector, or immunity genes that had one of these loss-of-function mutations (Table S3). Of the 251 decayed *aux* clusters, 70.5% had mutations in *vgrG*, 34.7% had mutated effectors, and 17.5% mutated immunity genes. Only 8.8% of *aux* clusters had mutations in the immunity gene while retaining evidently functional effectors and *vgrG* genes. Some *aux* types were more frequently decayed than others (Figure 4A, Figure S26, and Table S3). Clusters with nuclease effectors (*aux3, aux8, aux10, aux12, aux16, aux19, and aux21*) rarely decayed, as well as *aux13* and *aux23* whose effectors target unknown substrates. The most commonly decayed *aux* clusters were *aux1* (43/88 decayed), *aux2* (42/115 decayed) and *aux14* (36/75 decayed).

Most T6SS^+^ RSSC genomes have at least one decayed *aux* cluster. We investigated whether RSSC lineages varied in their proportion of decayed *aux* clusters, which could suggest that these lineages had less ecological pressure to maintain large toxin repertoires. As expected, the T6SS^null^ lineages (III, IV-BDB, and IV-SDC) mostly lack intact *aux* clusters although the BDB genomes had decayed *aux* clusters, and two T6SS^null^ phylotype III strains had intact *aux6* and *aux21* clusters (Figure S26, S29, and Table S3). Although phylotype IIB is T6SS^+^, IIB genomes encode fewer intact *aux* clusters than other T6SS^+^ clades, with a median of 5 intact *aux* clusters per genome (Figure 4B-C). A small clade within phylotype I that had five decayed *aux* clusters per genome (Figure 4, S26). Overall, there is no particularly strong phylogenetic pattern to the prevalence of decayed *aux* clusters.

### Mobile genetic elements facilitate horizontal acquisition of chromosomal *aux* clusters

The phylogenetic distribution of *aux* clusters among RSSC genomes suggests a complicated pattern of gene flow with frequent gain events in addition to the loss events documented above. We hypothesized that horizontal gene transfer (HGT) between RSSC clades may explain the convoluted phylogenetic pattern of *aux* cluster presence and absence. In bacteria, mobile genetic elements like phages and conjugative plasmids are common vehicles for the horizontal transmission of genes (67). To investigate if mobile genetic elements (MGEs) were associated with T6SS genes in RSSC genomes, we ran a combination of bioinformatic methods on the *aux* clusters, including genetic neighborhood visualization and comparison with Clinker (68), prophage prediction with Phaster (69) and domain analysis with NCBI CDD (60).

Of the 72 unique genetic neighborhoods around *aux* clusters, we classified 47.2% as MGE-associated, 44.4% as not MGE-associated, and the rest as inconclusive. We assigned the “inconclusive” status if there was minor but insufficient evidence that the cluster is associated with an MGE. For example, one cluster was encoded next to a single pseudogenized phage portal gene. We identified prophages (Myoviridae, Inoviridae, and Siphoviridae families), conjugative plasmids, and 16 other unclassified MGEs that were co-inherited with *aux* clusters (Figure 6A, Table S3). Of the prophages, the Myoviridae and Siphoviridae phages had highly conserved gene structure, whereas the filamentous Inoviridae phages were highly diverse.

**Figure 6.**
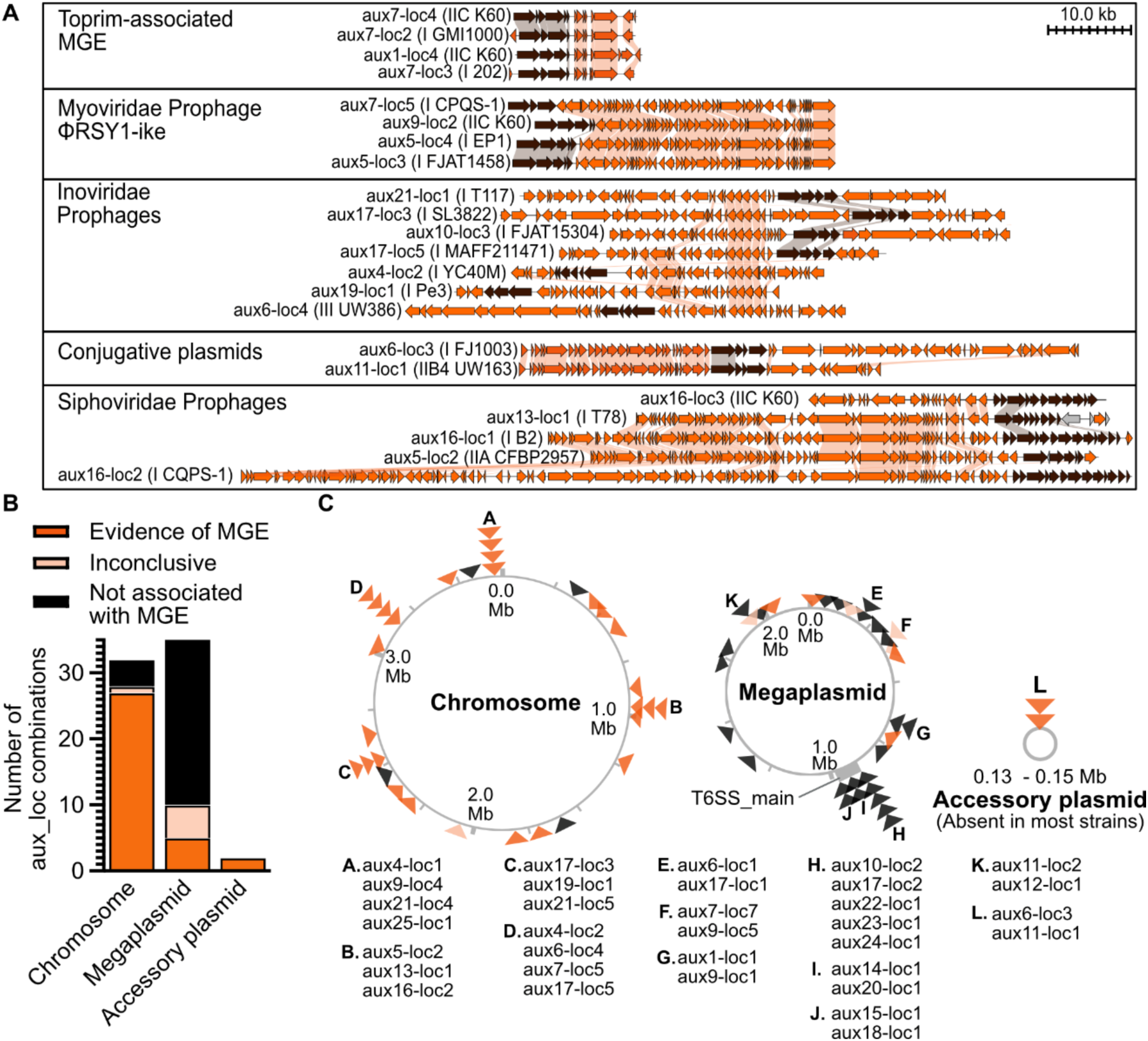
Mobile genetic elements mediate horizontal gene transfer of *aux* clusters. (**A**) Synteny analysis of representative *aux* loci associated with MGE. MGE genes are orange, *aux* cluster genes are brown, and all other genes are grey. (**B**) Proportion of of *aux* loci on the chromosomes, megaplasmids, and accessory plasmids that are MGE-associated, inconclusive, or not MGE-associated. (**C**) Specific locations of each *aux*-loci across the RSSC genomes. Each triangle represents one *aux* locus, and colors correspond to the legend in B. The letters identify hotspots where multiple *aux*-loci were found across the RSSC pangenome.

Many of the MGEs were associated with lipase *aux* types. The ϕRSY1-like prophages in the Myoviridae family carried certain lipase clusters: *aux5* at 3/6 locations, *aux7*, and *aux9* at 2/6 locations (70). Siphoviridae prophages carried a lipase cluster and two clusters with unknown targets: *aux5*, *aux13*, and *aux16* at 3/3 locations. An unclassified 8-gene MGE with a Toprim topoisomerase carried two lipase clusters *aux1* and *aux7* at 3/8 locations. Finally, the conjugative plasmids carried two lipase clusters *aux6* and *aux11*. Inoviridae prophages carried clusters with diverse toxins *aux4, aux6, aux10, aux17* at 2/5 locations*, aux19,* and *aux21*.

Several of the unclassified MGE contain anti-phage defense systems, such as a type 2 bacteriophage exclusion (BREX) system with the *aux21^loc4^* MGE and as DISARM restriction-modification system with the *aux6^loc4^* MGE (71, 72).

We investigated the genomic locations of MGE-associated, inconclusive, and non-MGE *aux* clusters. Surprisingly, we found that there was a strong linkage between chromosomal location and MGE- association (Figure 6B). Whereas 27/32 chromosomal *aux* clusters were MGE-associated, only 5/38 megaplasmid *aux* clusters were MGE-associated. All accessory plasmids with aux clusters were conjugative plasmid MGEs. We identified several hotspots on chromosome where *aux*-carrying MGEs inserted, and most of the hotspots were dominated by a single prophage family (Figure 6C). For example, hotspot B was dominated by the Siphoviridae phages, while C and D were both dominated by Inoviridae prophages. In contrast, hotspot A was occupied by four distinct unclassified MGEs.

## Discussion

Bacteria in the RSSC are aggressive pathogens that cause lethal wilt disease in plants, resulting in drastic losses of economically important crops. RSSC are renowned for manipulating a broad range of host plants with massive repertoires of 60-80 type 3 secreted effectors (73). Here, we document that RSSC are also evolutionary enriched in a large arsenal of type 6 secreted toxins. The high prevalence of T6SS genes in RSSC genomes suggests that possessing a T6SS is adaptive for the lifestyle of these pathogens. RSSC pathogens pass through a variety of microbial ecosystems to complete their disease cycle. When transmitting and infecting plants, they encounter a variety of ecological interactors. We speculate that the RSSC have an ecological role similar to spotted hyenas. Similar to how hyenas attack with their teeth and claws, RSSC bacteria may wield their T6SS against competitors (other RSSC and plant-colonizing bacteria), predators (bacterivorous amoebae) and, possibly, their prey (plant hosts).

Our data support the model that there is eco-evolutionary feedback between bacterial lifestyle and enrichment of T6SS-reated genes (26). We discovered that the RSSC and the *Acidovorax citrulli* species complex, two taxa of ecologically similar plant pathogens, have convergently evolved to deploy a large array of effectors from a single T6SS. These pathogens both cause acute infections of the plant xylem (74, 75). In contrast, plant symbiotic *Rhizobia* and *Agrobacterium* cause chronic infections with a spectrum of mutualistic to pathogenic outcomes (76–78). These chronic symbionts have only a single T6SS and only a handful of effectors, which suggests a reduced need to combat diverse microbes and a greater emphasis on reducing collateral damage to the host. In contrast, other Burkholderiaceae encode up to 6 distinct T6SSs in their genomes (79, 80), which is consistent with a model that organisms with complex lifestyles benefit from having multiple T6SSs that are independently regulated and specialized for different targets and contexts.

The relationship between bacterial lifestyle and T6SS gene content is also evident within the RSSC. Most RSSC pathogens are transmitted through soil or fresh water. However, two of the phylotype IV clades that lack a T6SS have adopted novel lifestyles: the strains causing Sumatra Disease of Clove (SDC) and Blood Disease of Banana (BDB) (81, 82). SDC strains are spread by piercing sucking insects (83) while BDB is mechanically transmitted by insects or agricultural tools (84). While the two SDC-lineage genomes lacked *aux* clusters, they encoded two T6SS core genes (*gh10* and *ompA*). Similarly, all three genomes in the BDB lineage lacked T6SS core genes but contained pseudogenized *aux* clusters. Overall, this pattern suggests that the T6SS is ancestral to the RSSC and has been recently lost in T6SS^null^ lineages. Like many bacteria that transition to a host-restricted lifestyle, these lineages have undergone a reduction in genome size. While SDC and BDB strains have chromosomes of comparable size to soil-transmitted relatives, their megaplasmids are reduced in size (85).

Although the T6SS is widely conserved in soil-transmitted RSSC, the absence of the T6SS in several soil-transmitted lineages demonstrates that the T6SS is non-essential for this lifestyle. Soil-transmitted RSSC that lack a T6SS included all but one of the phylotype III genomes and a minor clade of phylotype I (Figure 2C). Three of the T6SS^null^ phylotype III strains had intact *aux* clusters, which could suggest that the T6SS was recently lost from this lineage. However, two of the three intact *aux* clusters were associated with mobile genetic elements (MGEs), so it is alternatively possible these *aux* clusters were recently gained through HGT after the loss of the T6SS.

Within phylotype II, several RSSC lineages cause Moko disease of bananas and plantains. Pioneering studies on Moko disease by Luis Sequeira and Ivan Buddenhagan demonstrated that the causal pathogens are facultatively transmitted by either soil transmission or contact transmission of insects with sap, similar to BDB (86, 87). We and other groups have sequenced isolates from the Sequiera and Buddenhagan collection from the 1960s Moko epidemic (88–91), and we now know that multiple RSSC lineages were responsible for the epidemic, including T6SS^+^ lineages (IIB-4 and IIB-3) and T6SS^null^ lineages (IIA-6). Further epidemiological research is needed to understand how variation in transmission routes shapes the evolution and behavior of RSSC lineages.

Our results provide new insight into the epidemiology of RSSC in regions with multiple lineages. It has long been known that RSSC strains can inhibit each other’s growth and competitively exclude each other *in planta* (92, 93). A thorough epidemiological survey of Malagasy vegetable plots demonstrated that T6SS^+^ phylotype I strains are displacing the T6SS^null^ phylotype III strains native to the island (94). Subsequent functional analysis demonstrated that the phylotype I Malagasy strains secrete bacteriocin toxins that inhibit growth of the phylotype III strains (95). Our results suggest that the T6SS may confer an additional advantage to phylotype I strains when directly competing against T6SS^null^ phylotype III strains *in planta*.

The convoluted phylogenetic distribution of *aux* clusters suggests that RSSC populations have dynamic T6SS gene flow with frequent gain and loss events. Although there is evidence that certain *aux* types have been primarily vertically inherited, most *aux* clusters are clearly horizontally inherited based on their phylogenetic distribution patterns and the context of their genetic neighborhoods. Notably, the ϕRSY1-like Myoviridae prophages definitively transfer *aux* clusters. ϕRSY1 was originally from soil from a *Ralstonia*-infested field, and whole genome sequencing of the purified virion particles confirms that the genomes of ϕRSY1-like phages encode *aux* clusters and, thus, can transmit *aux* clusters.

Like many bacteria in the Burkholderiaceae, RSSC have a bipartite genome. When bacteria have a secondary replicon like the megaplasmid, the genes on the secondary replicon evolve more quickly than chromosomal genes (64–66). Intriguingly, we discovered that *aux* clusters are dramatically enriched on the megaplasmid. We speculate that the enrichment of *aux* clusters on the evolutionarily dynamic megaplasmid could allow RSSC lineages to rapidly diversify with respect to their intraspecies and interspecies competitive behavior. In-depth studies on the molecular evolution of *aux* cluster genes are needed to understand if they evolve at different rates than other RSSC genes and if there are differences in the evolutionary behavior of megaplasmid and chromosomal MGE-associated *aux* clusters.

In *Ralstonia*’s bipartite genome, we found clear distinctions in the mechanisms for horizontal gene transfer for *aux* clusters on the chromosome vs. megaplasmid. Chromosomal *aux* clusters were predominantly MGE-associated, which is consistent with prior reports that RSSC prophages are moderately enriched on the evolutionarily stable chromosome and that they target specific insertion sequences (96). In contrast, megaplasmid *aux* clusters were rarely associated with MGEs. Nevertheless, the patterns of phylogenetic distribution suggest that both chromosomal and megaplasmid *aux* clusters have been horizontally transmitted. This opens the question—what genetic mechanisms contributed to horizontal gene flow of megaplasmid *aux* clusters? RSSC are naturally competent (97), so they could readily acquire genes by uptake of environmental DNA and integration by homologous recombination. However, it is surprising that there would be a bias towards homologous recombination for the megaplasmid.

The presence of *aux* clusters in the genomes of phages indicates that these phages may function as mutualists of RSSC. Across the diversity of RSSC-infective phages, most do not transport *aux* clusters (96, 98–100). Nevertheless, with growing interest in employing phages for control of bacterial plant pathogens (99, 101–105), it will remain important to evaluate candidate biocontrol agents for their ability to improve the ecological fitness of the pathogens.

In closing, we propose that evolution has positioned the RSSC to be an ideal model to understand the role of the T6SS in shaping pathogen populations and host-associated ecosystems. Our systematic analysis opens a plethora of evolutionarily grounded questions for future investigation. For example, do RSSC pathogens target novel cellular targets with their nine effector families that lack known toxin domains? Are there genetic and epigenetic mechanisms that promote preferential integration of *aux* clusters on the megaplasmid? Moreover, what other physiological functions are enriched on the megaplasmid and secondary replicons in other bacteria with multipartite genomes? RSSC genomes also encode large repertoires of other genes that allow them to sense and change their environments, including root-sensing MCP chemoreceptors (106), plant-manipulating T3SS effectors (73), and anti-phage defense systems (107). Do these genes exhibit biased distribution across the replicons? We anticipate that this study will fuel many new discoveries in T6SS biology and pathogen evolution.

## Materials and Methods

### Identification of genes encoding the type 6 secretion system (T6SS) machinery in RSSC genomes

All publicly available genomes in the genus *Ralstonia*, including *R. syzygii* genomes deposited as “Blood Disease Bacterium” strains, were downloaded from NCBI and uploaded to KBase (108) for analysis. Genomes were analyzed with the Genome Taxonomy Database Toolkit GTDB-Tk - *v1.7.0* to identify the genomospecies (34–39). We used CheckM - *v1.0.18* to evaluate the completeness and contamination of the assemblies. Only genomes with completeness greater than 99.82 % and contamination less than 0.96 % were retained for further analysis. We also excluded assemblies with 100 or more contigs. With the selected assemblies we generated a phylogenetic tree of RSSC using Insert Genome Into SpeciesTree *– v.2.2.0.* The KBase SpeciesTree uses 49 genes broadly conserved across bacteria to build the phylogenetic tree. The protein sequence for each GMI1000 T6SS component was queried against the *Ralstonia* genomes using BLASTp - *v2.13.0*. We considered all BLASTp results with ≥ 20% identity and ≥ 80% coverage to be hits. We analyzed phylogenetic patterns of T6SS gene presence or absence in the RSSC phylogenetic tree on iTOL (109).

We selected high quality genomes for detailed analysis of the repertoires of *vgrG*-linked effector/immunity gene clusters. Limiting the analysis to complete genomes would have excluded most phylotype II, III, and IV genomes, so we included any genome assembled into as many as 28 contigs. We carried out a series of low stringency BLASTp searches against the RSSC genomes with VgrG protein sequences from phyl. I GMI1000, phyl. II IBSBF1503, and phyl. IV PSI07 (Parameters: ≥ 1% identity, ≥ 1% coverage, and Bit Score ≥ 10). Vgr BLASTp results were exported and further analyzed in Excel.

### Classifying the Vgr-linked effector/immunity gene clusters through synteny analysis

We downloaded the genetic neighborhood of each *vgrG* gene from NCBI. The gene locus was queried against the NCBI Nucleotide database. From the corresponding Graphics page, we downloaded a Genbank Flat file that encompassed a 10-200 kb region surrounding the *vgrG* gene. The .flat files were converted to .gbk, and the synteny analysis and visualization tool Clinker was iteratively used to identify gene clusters with shared genetic architecture (68). *vgrG*-linked clusters were named *aux1-aux25,* which is short for auxiliary cluster because these effector/immunity clusters are not core T6SS genes. Per genome, the set of *vgrG*-linked clusters were matched to references for each auxiliary cluster (*aux1- aux25*) using Clinker (68).

Because the VgrG BLASTp search would not identify *aux* clusters with pseudogenized or missing *vgrG* genes, we carried out further BLASTp queries with the effector and immunity protein sequences. We exported the gene neighborhoods from NCBI and used Clinker to confirm the *aux-*type of each cluster.

### Identification of T6SS and *vgrG* genes in representative Burkholderiaceae genomes

The Genome Taxonomy Database (GTDB) (34–39) was used to identify complete genomes in the Burkholderiaceae family. GTDB classifies genomes into genomospecies based on a 95% ANI threshold. Per GTDB species, we selected one representative genome to import into a public KBase Narrative (https://narrative.kbase.us/narrative/142785). We used BLASTp as described above to identify T6SS_main genes and VgrG proteins. The full results are present in Table S1.

### Classifying the genetic architecture of T6SS_main loci across the *Ralstonia* genus

Clinker was used to visually compare the genetic architectures of T6SS_main loci. We downloaded the genetic neighborhood of each T6SS_main locus from NCBI and compared them to reference T6SS_main loci from plant-colonizing bacteria (49, 110). The phylogenetic distributions of the T6SS subtypes were visualized with iTol (109).

### Identifying effectors, immunity proteins, and adaptors encoded in each *aux* cluster

We queried protein sequences (“WP_” accessions) against the literature using PaperBlast (61) and the NCBI Conserved Domain Database (CDD) (60) using CD-Search. Effectors were identified based on homology with bona fide T6SS effectors (e.g., PaperBlast matches to Tle1, Tle3, Tle4, and Tle5 phospholipases). Immunity proteins were identified based on homology to bona fide immunity proteins (PaperBlast) and/or the presence of known immunity domains. Adapters were identified based on the presence of domains like DUF1795, DUF2169, or DUF4123. Sometimes the VgrG or effector encoded a PAAR spike motif and sometimes the cluster had a small, standalone PAAR-containing protein.

### Identifying genetic decay in the clusters and orphan immunity genes

We recorded which genes were annotated as pseudogenized. Moreover, the Clinker visualizations enabled us to identify transposons and/or insertion sequences that had been inserted into and disrupted genes. The visualizations revealed if an effector or *vgrG* gene was conspicuously absent from a cluster.

### Identifying the genomic location of each *vgrG*-linked *aux* cluster

Clinker was used to identify which *aux* clusters were in the same genomic location across genomes. Per *aux*-type (i.e., all “*aux1*” clusters), Clinker was run, and unique auxiliary loci were identified and named. Each auxiliary locus was mapped to its relative position with respect to the GMI1000 genome by BLAST-searching neighboring genes against the GMI1000 genome and retrieving a Genbank Flat file of the approximate region. Auxiliary loci and their mapped location are referred to as “*aux1-loc1*” and “*aux1- loc2*”. Synteny between the GMI1000 sequence and the *aux* cluster’s neighborhood was visualized with Clinker. If GMI1000 encoded the exact cluster, the locus for the *vgrG* gene was recorded. If GMI1000 lacked the cluster, the nearest GMI1000 gene upstream of the *vgrG* was recorded.

When downloading Genbank Flat files from complete genomes, we recorded whether the cluster was encoded on the chromosome, megaplasmid, or a small accessory plasmid. For incomplete genomes, we inferred whether the cluster was on the chromosome or megaplasmid by matching the cluster’s *aux- lo*c to a syntenous cluster from a complete genome.

To visualize the relative location of each cluster across the replicons (Figure S4-S24), we used Clinker to compare a representative *aux* cluster from each of the 90 *aux-loc* subtypes to the relative region in GMI1000. The GMI1000 gene upstream of each VgrG was recorded as the “nearby location marker” in Table S3.

### Identifying *aux* clusters diversity in RSCC

To assess *aux* clusters diversity in RSSC we measured Bray-Curtis dissimilarity matrix which accounts for the presence, absence, and relative abundance of *aux* clusters. We used ‘vegdigest’ of the ‘vegan v.2.6-2’ package in RStudio (version 2023.03.0+386) to compute the Bray-Curtis dissimilarity matrix. With this matrix, we constructed an UPGMA tree with ‘hclust’ of the ‘Phangorn v.2.11.1’ in RStudio (version 2023.03.0+386).

### Identifying mobile genetic elements associated with *aux* clusters

To identify MGEs associated with various T6SS auxiliary clusters, we started with BLASTp searches on the *aux* clusters with ∼100 Kb on each side of their genetic neighborhoods in PHASTER identification and annotation of prophage tool (69). In parallel, we checked gene annotations of the genetic neighborhoods. To check if hypothetical proteins could be associated to mobile genetic elements, we checked for the presence of any mobile genetic elements-related domains on the Conserved Domain Database of NCBI (60). Based on this analysis we recorded the presence, inconclusive, or absence of MGE on the chromosome, megaplasmid, and the conjugative plasmid (when present). The barplot showing the proportions of the presence, inconclusive, or absence of MGE (Figure 6) was built using GraphPad Prsim (version 10.0.1). We highlighted the *aux* clusters associated with MGE in orange using AFFINITY Designer (version 1.10.6). *aux* clusters and corresponding locations were summarized on the chromosome, megaplasmid, and conjugative plasmid maps using AFFINITY Designer (version 1.10.6). The synteny

## Acknowledgments

We would like to thank Dr. Jonathan Jacobs for insightful recommendations about senteny analysis. We would also like to thank Dr. Anneliek ter Horst, Dr. Joanne Emerson, and Dr. Poliane Alfenas-Zerbini for sharing expertise in phage genomics.

