## Supplementary information for "A pangenomic atlas reveals that eco-evolutionary dynamics shape plant pathogen type VI secretion systems"

**Table S1. BLASTp results of Burkholderiaceae species T6SS genes**
**Table S2. BLASTp results of RSSC species T6SS genes**
**Table S3. Summary of *aux* clusters and associated-MGE**
**Table S4. Bray-Curtis dissimilarity matrix of the complete *aux* clusters**

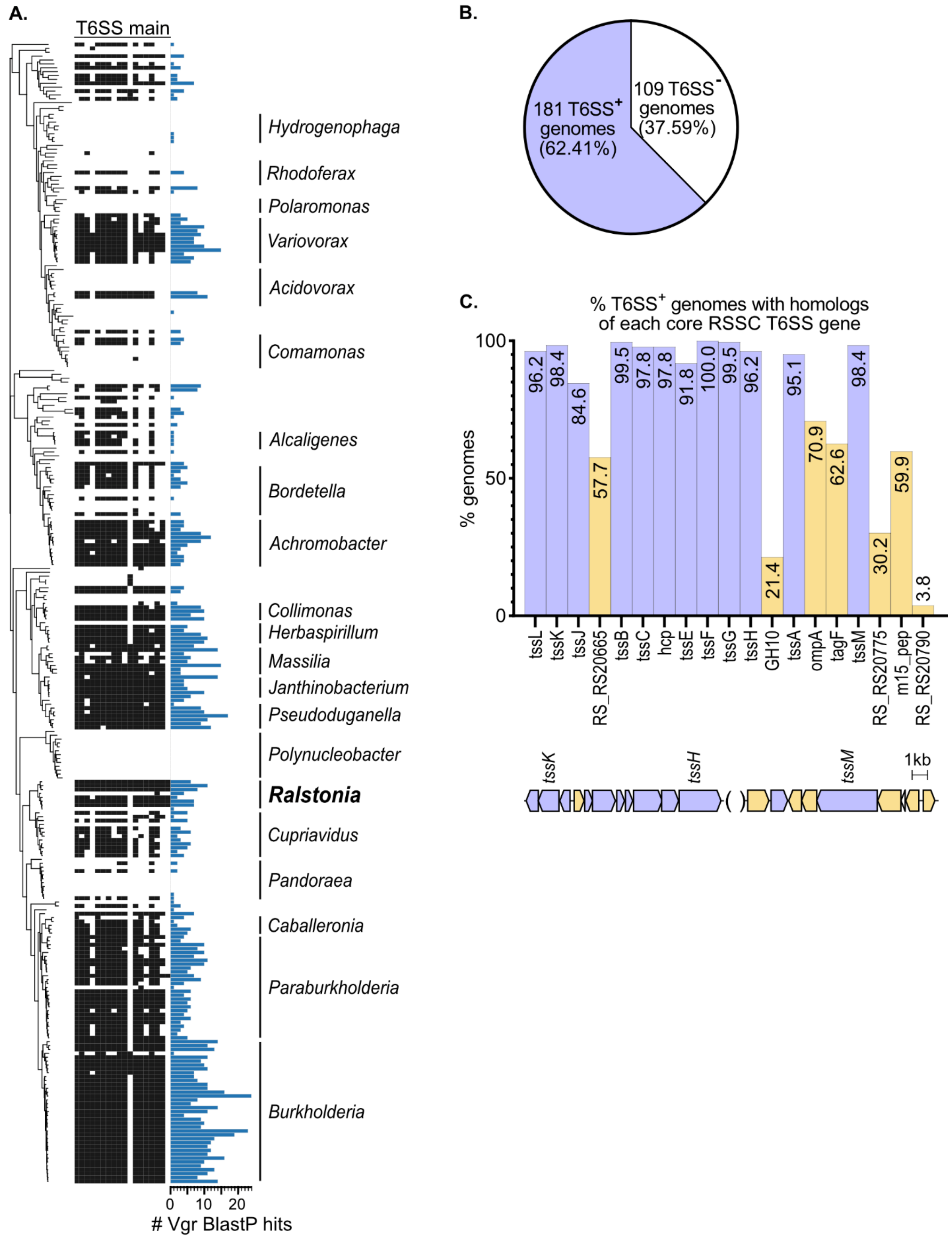


**Figure S1. Diverse Burkholderiaceae species representatives encode some, but not all, of the conserved RSSC T6SS_main genes.** We investigated the prevalence of T6SS secretion system genes across diverse

Burkholderiaceae. Using the Genome Taxonomy Database (GTDB), we identified complete genomes that represent 290 genomospecies within the Burkholderiaceae family. We built a custom database of these genomes in KBase and carried out BlastP searches of 18 genes that all T6SS+ RSSC strains encode in the main locus and estimated Vgr copy number with BlastP. (A) Phylogenetic distribution of BlastP hits for presence/absence of T6SS genes and copy number of Vgr alleles. The order of the T6SS genes is the same as the barchart in C. Genera with many species representatives are labeled. A PDF of the full tree is available on FigShare: https://doi.org/10.6084/m9.figshare.23499141.v1. (B) The proportion of the 290 genomes that encode at least one T6SS. (C) The prevalence of RSSC's core T6SS genes across Burkholderiaceae genomes. Blue indicates genes identified in most T6SS+ genomes and yellow indicates genes that are more rarely encoded. The conserved and variable T6SS genes are enriched in different clusters of the RSSC T6SS_main.

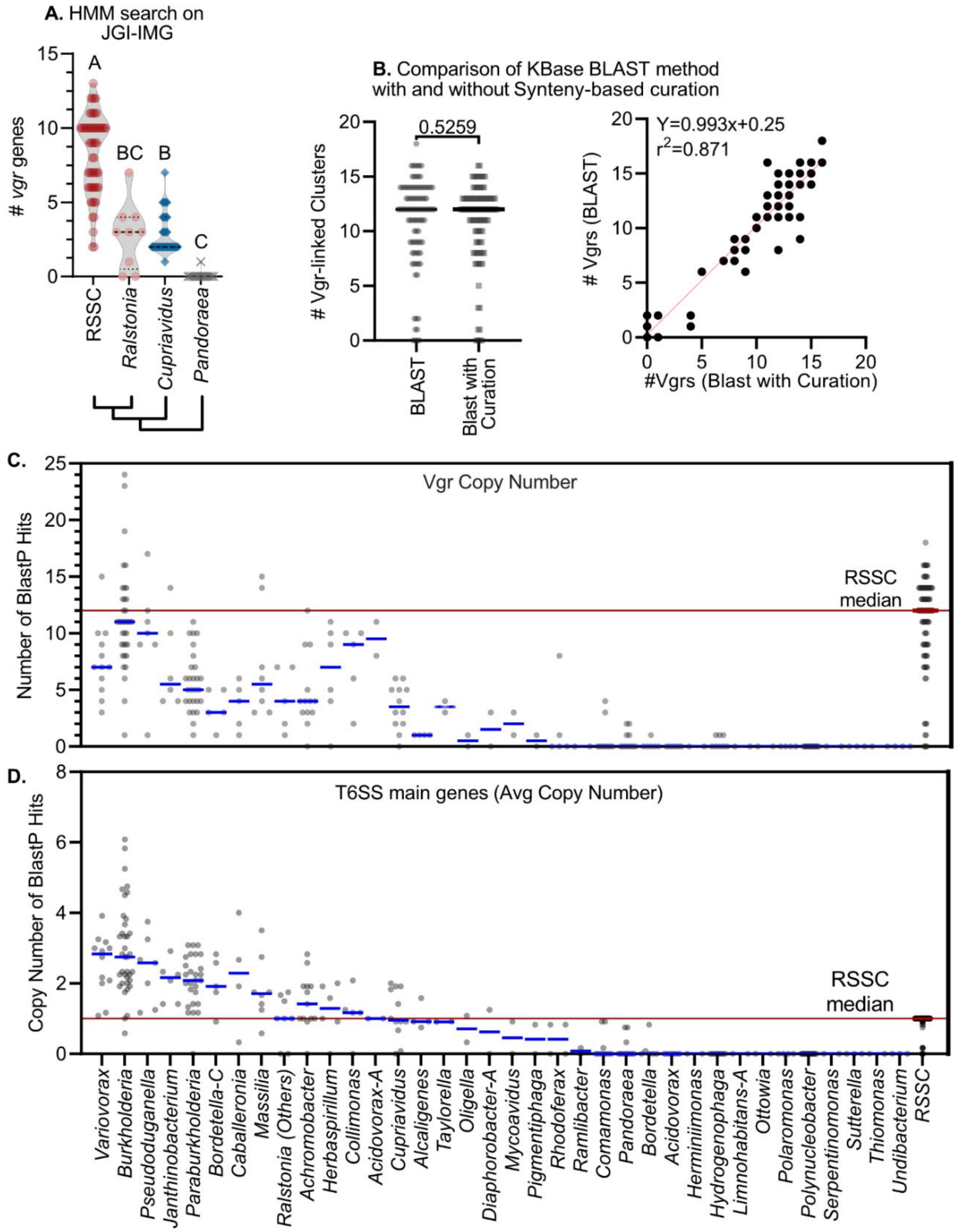


**Figure S2. Additional Data showing RSSC genomes are enriched in Vgr-linked toxin/anti-toxin clusters.** We quantified Vgr abundance in RSSC genomes with three independent methods: (**A-B**) HMM-based searches with TIGR03361 on the JGI IMG platform, BlastP-based searches with 6 distinct Vgr alleles using custom genome databased on KBase, and curation of the BlastP search results for high quality RSSC genomes using Clinker synteny analysis for regions surrounding Blast results for Vgr, toxin, or antitoxin alleles. (**B**) The results of the BlastP-only approach were similar to results of the BlastP-with-curation approach. The approaches yield statistically insignificant results based on the Mann-Whitney test. A linear regression between the two approaches is displayed on the XY plot (y=0.993x+0.25 with $r^2 = 0.871$). (**C-D**) Comparison of T6SS gene content across Burkholderiaceae genomes corresponding to Fig 1A. For the T6SS_main, each circle represents a single genome's average number of BlastP hits for the Core T6SS components: TssA, TssB, TssC, TssE, TssF, TssG, TssH, TssJ, TssK, TssL, TssM, and Hcp.

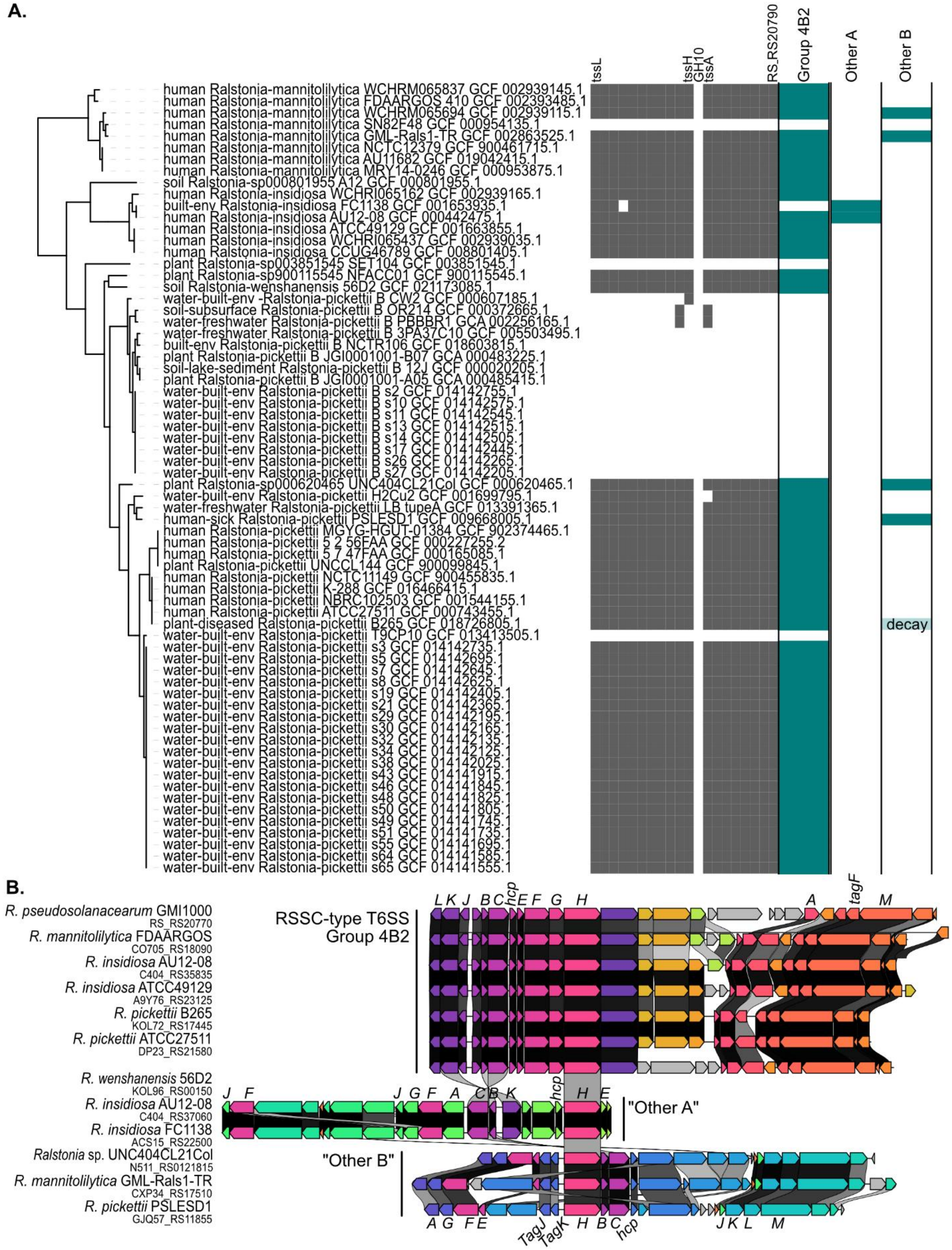
A.
tssL
tssH
GH10
tssA
RS_RS20790
Group 4B2
Other A
Other B
human Ralstonia-mannitolilytica WCHRM065837 GCF 002939145.1
human Ralstonia-mannitolilytica FDAARGOS 410 GCF 002393485.1
human Ralstonia-mannitolilytica WCHRM065694 GCF 002939115.1
human Ralstonia-mannitolilytica SN82F48 GCF 000954135.1
human Ralstonia-mannitolilytica GML-Rals1-TR GCF 002863525.1
human Ralstonia-mannitolilytica NCTC12379 GCF 900461715.1
human Ralstonia-mannitolilytica AU11682 GCF 019042415.1
human Ralstonia-mannitolilytica MRY14-0246 GCF 000953875.1
soil Ralstonia-sp000801955 A12 GCF 000801955.1
human Ralstonia-insidiosa WCHRI065162 GCF 002939165.1
built-env Ralstonia-insidiosa FC1138 GCF 001653935.1
human Ralstonia-insidiosa AU12-08 GCF 000442475.1
human Ralstonia-insidiosa ATCC49129 GCF 001663855.1
human Ralstonia-insidiosa WCHRI065437 GCF 002939035.1
human Ralstonia-insidiosa CCUG46789 GCF 008801405.1
plant Ralstonia-sp003851545 SET104 GCF 003851545.1
plant Ralstonia-sp900115545 NFACC01 GCF 900115545.1
soil Ralstonia-wenshanensis 56D2 GCF 021173085.1
water-built-env -Ralstonia-pickettii B CW2 GCF 000607185.1
soil-subsurface Ralstonia-pickettii B OR214 GCF 000372665.1
water-freshwater Ralstonia-pickettii B PBBBR1 GCA 002256165.1
water-freshwater Ralstonia-pickettii B 3PA37C10 GCF 005503495.1
built-env Ralstonia-pickettii B NCTR106 GCF 018603815.1
plant Ralstonia-pickettii B JGI0001001-B07 GCA 000483225.1
soil-lake-sediment Ralstonia-pickettii B 12J GCF 000020205.1
plant Ralstonia-pickettii B JGI0001001-A05 GCA 000485415.1
water-built-env Ralstonia-pickettii B s2 GCF 014142755.1
water-built-env Ralstonia-pickettii B s10 GCF 014142575.1
water-built-env Ralstonia-pickettii B s11 GCF 014142545.1
water-built-env Ralstonia-pickettii B s13 GCF 014142515.1
water-built-env Ralstonia-pickettii B s14 GCF 014142505.1
water-built-env Ralstonia-pickettii B s17 GCF 014142445.1
water-built-env Ralstonia-pickettii B s26 GCF 014142265.1
water-built-env Ralstonia-pickettii B s27 GCF 014142205.1
plant Ralstonia-sp000620465 UNC404CL21Col GCF 000620465.1
water-built-env Ralstonia-pickettii H2Cu2 GCF 001699795.1
water-freshwater Ralstonia-pickettii LB tupeA GCF 013391365.1
human-sick Ralstonia-pickettii PSLESD1 GCF 009668005.1
human Ralstonia-pickettii MGYG-HGUT-01384 GCF 902374465.1
human Ralstonia-pickettii 5 2 56FAA GCF 000227255.2
human Ralstonia-pickettii 5 7 47FAA GCF 000165085.1
plant Ralstonia-pickettii UNCCL144 GCF 900099845.1
human Ralstonia-pickettii NCTC11149 GCF 900455835.1
human Ralstonia-pickettii K-288 GCF 016466415.1
human Ralstonia-pickettii NBRC102503 GCF 001544155.1
human Ralstonia-pickettii ATCC27511 GCF 000743455.1
plant-diseased Ralstonia-pickettii B265 GCF 018726805.1
water-built-env Ralstonia-pickettii T9CP10 GCF 013413505.1
water-built-env Ralstonia-pickettii s3 GCF 014142735.1
water-built-env Ralstonia-pickettii s5 GCF 014142695.1
water-built-env Ralstonia-pickettii s7 GCF 014142645.1
water-built-env Ralstonia-pickettii s8 GCF 014142625.1
water-built-env Ralstonia-pickettii s19 GCF 014142405.1
water-built-env Ralstonia-pickettii s21 GCF 014142365.1
water-built-env Ralstonia-pickettii s29 GCF 014142195.1
water-built-env Ralstonia-pickettii s30 GCF 014142165.1
water-built-env Ralstonia-pickettii s32 GCF 014142135.1
water-built-env Ralstonia-pickettii s34 GCF 014142125.1
water-built-env Ralstonia-pickettii s38 GCF 014142025.1
water-built-env Ralstonia-pickettii s43 GCF 014141915.1
water-built-env Ralstonia-pickettii s46 GCF 014141845.1
water-built-env Ralstonia-pickettii s48 GCF 014141825.1
water-built-env Ralstonia-pickettii s50 GCF 014141805.1
water-built-env Ralstonia-pickettii s49 GCF 014141745.1
water-built-env Ralstonia-pickettii s51 GCF 014141735.1
water-built-env Ralstonia-pickettii s55 GCF 014141695.1
water-built-env Ralstonia-pickettii s64 GCF 014141585.1
water-built-env Ralstonia-pickettii s65 GCF 014141555.1
decay
B.
R. pseudosolanacearum GMI1000
RS_RS20770
R. mannitolilytica FDAARGOS
CO705_RS18090
R. insidiosa AU12-08
C404_RS35835
R. insidiosa ATCC49129
A9Y76_RS23125
R. pickettii B265
KOL72_RS17445
R. pickettii ATCC27511
DP23_RS21580
R. wenshanensis 56D2
KOL96_RS00150
R. insidiosa AU12-08
C404_RS37060
R. insidiosa FC1138
ACS15_RS22500
Ralstonia sp. UNC404CL21Col
N511_RS0121815
R. mannitolilytica GML-Rals1-TR
CXP34_RS17510
R. pickettii PSLESD1
GJQ57_RS11855
RSSC-type T6SS
Group 4B2
L K J B C hcp E F G H
A tagF M
J F J G F A C B K hcp H E
"Other A"
"Other B"
A G F E TagJ TagK H B C hcp J K L M

**Figure S3. The RSSC T6SS is broadly conserved across the *Ralstonia* genus.**

(**A**) Left: phylogenetic tree of non-RSSC *Ralstonia* genomes, generated using the KBase SpeciesTree app. Middle: Grey squares show the presence of the RSSC T6SS_main genes across the non-RSSC *Ralstonia* genomes. Right: Teal rectangles show the presence of the RSSC-type T6SS (Group 4B2 according to Bernal et al. 20xxx) or two rare types of T6SS clusters. (B) Synteny of representative T6SS_main gene clusters from the three types.

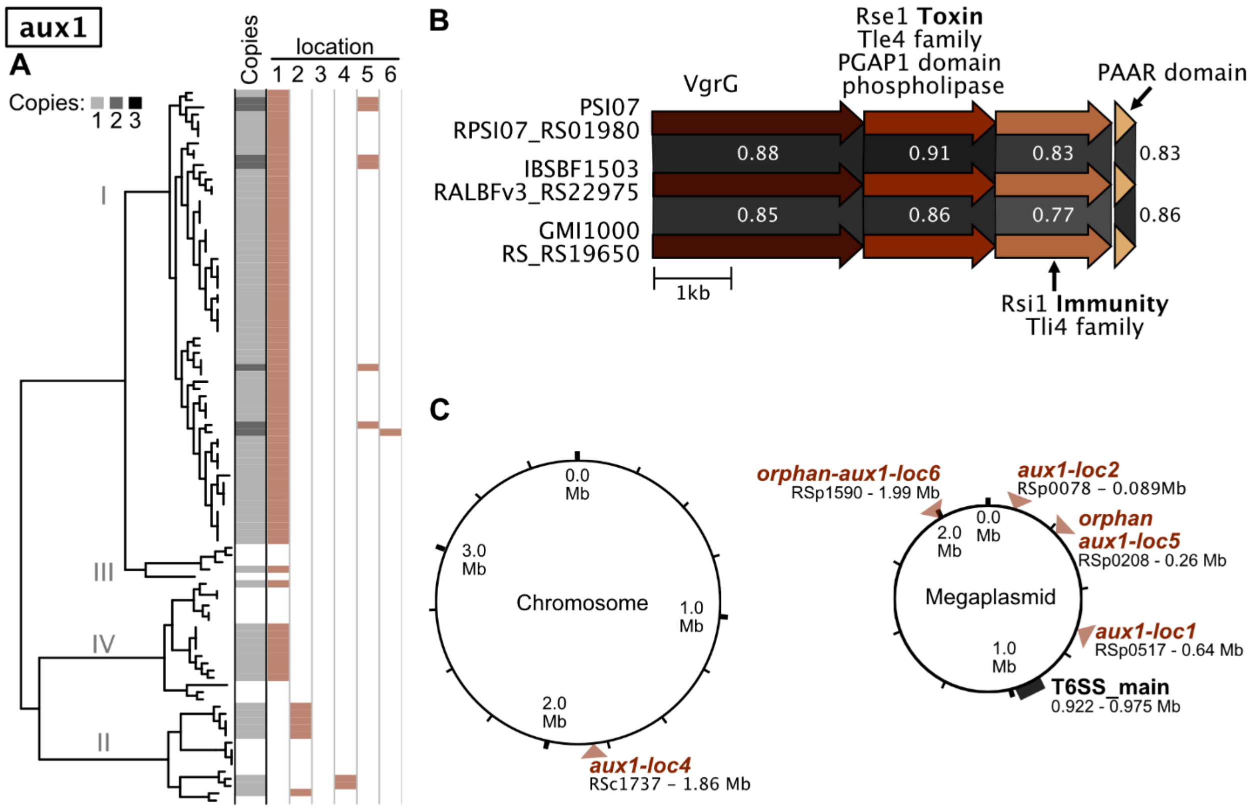


**Figure S4. Phylogenetic distribution, genetic organization/synteny, and chromosomal location of auxiliary *vgrG*-linked cluster 1 (*aux1*).** (**A**) Phylogenetic distribution and copy number of *aux1* across high-quality RSSC genomes. (**B**) Genetic organization/synteny of *aux1* from 3 genomes: phyl. I GMI1000, phyl. IIB-4 IBSBF1503, and IV-10 PSI07. *aux1* encodes a VgrG, a Tle4-family phospholipase with a PGAP1 domain, a Tli4 immunity protein, and a PAAR domain protein. Greyscale links indicate the amino acid identity between homologs. (**C**) *aux1* clusters were identified at six locations across the chromosome and megaplasmid, and these locations are shown relative to the GMI1000 replicons. Panel A indicates which genomes encode *aux1* at each location. The figure was generated with a combination of KBase BLASTp, iToL, Clinker, and Affinity Designer.

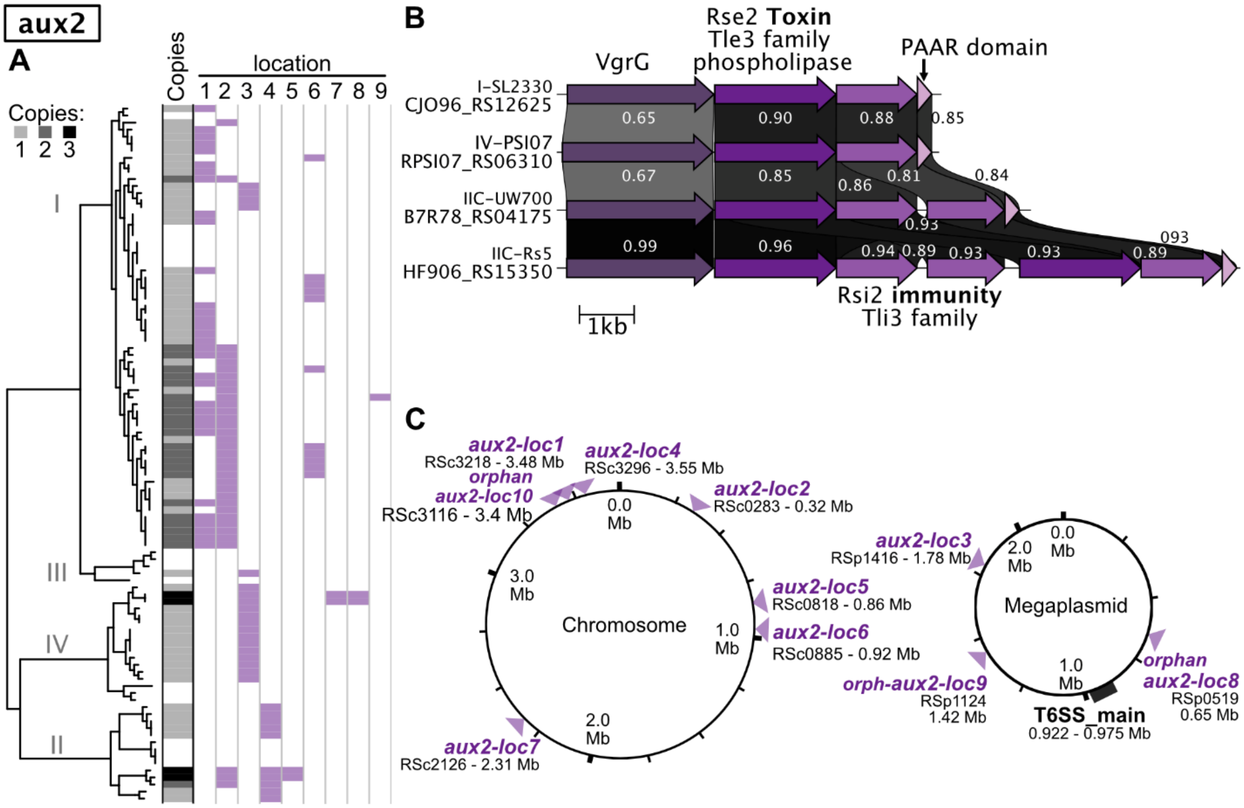


**Figure S5. Phylogenetic distribution, genetic organization/synteny, and chromosomal location of auxiliary *vgrG*-linked cluster 2 (*aux2*).** (**A**) Phylogenetic distribution and copy number of *aux2* across high-quality RSSC genomes. (**B**) Genetic organization/synteny of the cluster from 4 genomes: phyl. I SL2330, phyl. IIC-7 UW700, IIC-7 Rs5, and IV-10 PSI07. *aux2* encodes a VgrG, a Tle3-family phospholipase, one-or-more Tli3 immunity protein(s), and a PAAR domain protein. Greyscale links indicate the global amino acid identity between homologs. (**C**) *aux2* clusters were identified at nine locations across the chromosome and megaplasmid, and these locations are shown relative to the GMI1000 replicons. Panel A indicates which genomes encode the cluster at each location. The figure was generated with a combination of KBase BLASTp, iToL, Clinker, and Affinity Designer.

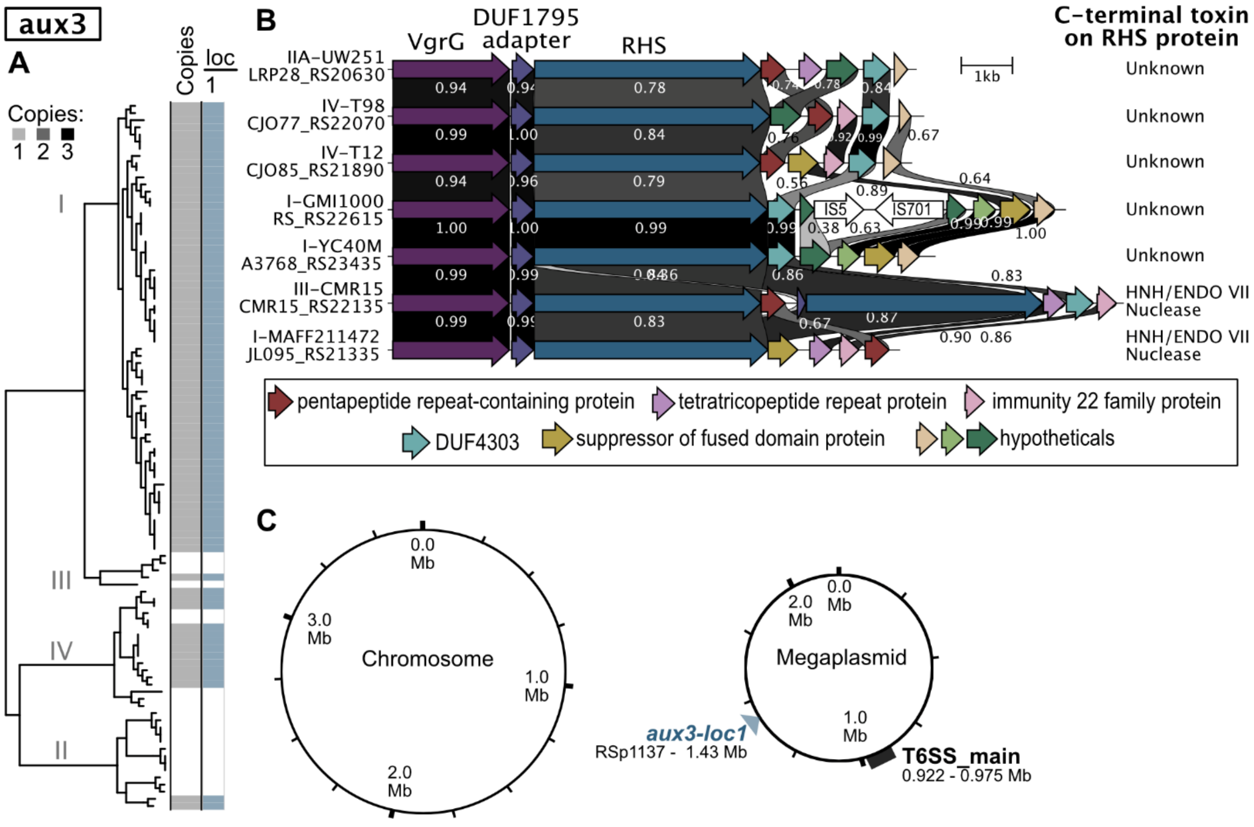


**Figure S6. Phylogenetic distribution, genetic organization/synteny, and chromosomal location of auxiliary *vgrG*-linked cluster 3 (*aux3*).** (**A**) Phylogenetic distribution and copy number of *aux3* across high-quality RSSC genomes. (**B**) Genetic organization/synteny of the cluster from 7 genomes: phyl. I GMI1000, phyl. I MAFF211472, phyl. I YC40M, phyl. IIA UW251, phyl. III CMR15, phyl. IV T98, and phyl. IV T12. *aux3* encodes a VgrG, a DUF1795 adapter protein, an RHS with a putative C-terminal effector, and four-or-more variable proteins: a pentapeptide repeat-containing protein, a tetratricopeptide repeat protein, and imm22 family immunity protein, a DUF4303, a suppressor of fused domain protein, and hypothetical(s). Some of the C-terminal effectors are putative HNH/ENDO VII nucleases while others do not encode known domains. The small genes downstream of the RHS effector are likely an array of immunity proteins. Greyscale links indicate the global amino acid identity between homologs. (**C**) *aux3* clusters were identified at one location on the megaplasmid, and the location is shown relative to the GMI1000 replicons. The figure was generated with a combination of KBase BLASTp, iToL, Clinker, and Affinity Designer.

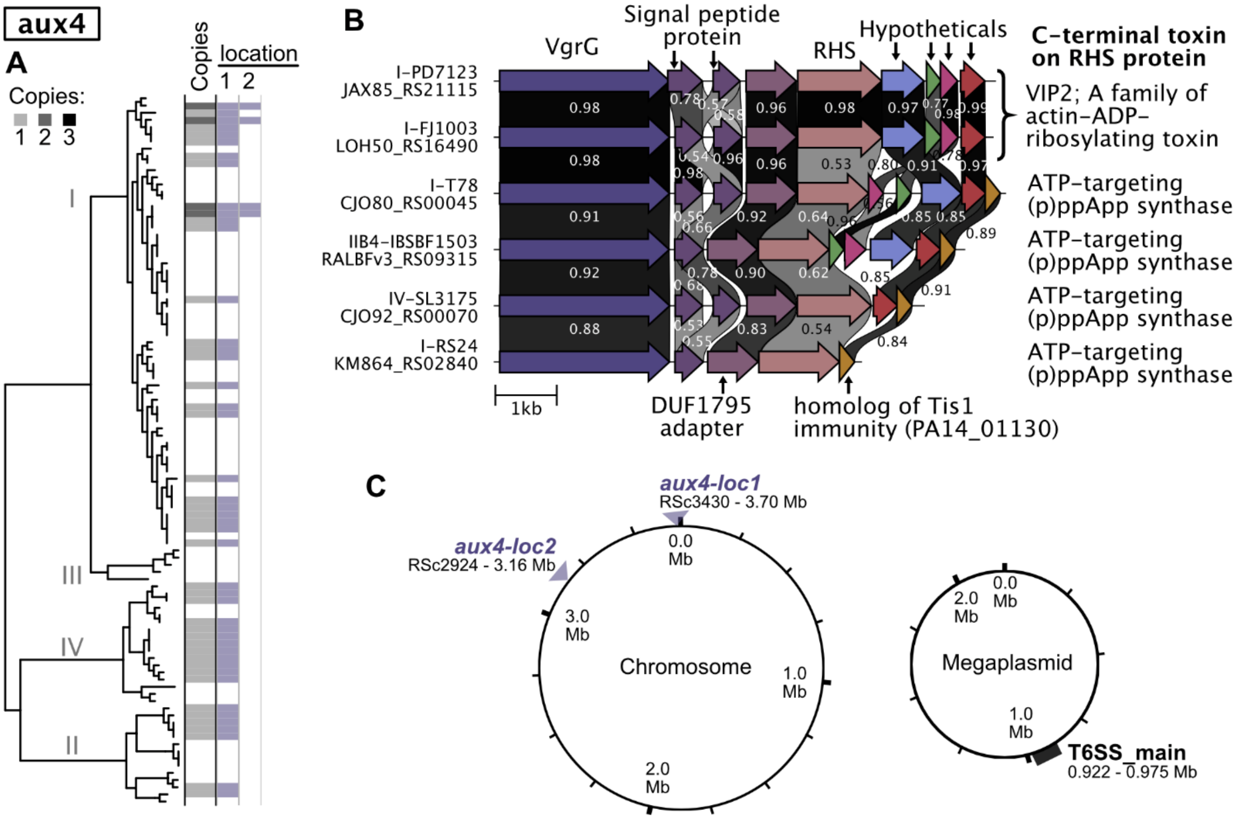


**Figure S7. Phylogenetic distribution, genetic organization/synteny, and chromosomal location of auxiliary *vgrG*-linked cluster 4 (*aux4*).** (**A**) Phylogenetic distribution and copy number of *aux4* across high-quality RSSC genomes. (**B**) Genetic organization/synteny of the cluster from 6 genomes: phyl. I PD7123, phyl. I FJ1003, phyl. I T78, phyl. I RS24, phyl. IIB-4 IBSBF1503, and IV SL3175. *aux4* encodes a VgrG, one-or-more signal peptide proteins, a DUF1795 adapter protein, an RHS with a putative C-terminal effector, and one-or-more variable proteins: a homolog of the *P. aeruginosa* Tis1 immunity protein or four distinct hypothetical(s). The small genes downstream of the RHS effector likely encode an array of immunity proteins. Some of the C-terminal effectors are annotated as VIP2 actin-ADP-ribosylating effectors while others have similarity to the ATP-targeting (p)ppApp synthases Tas1 (Ahmad et al., 2019), including conserved active site motifs and the catalytic glutamate. Greyscale links indicate the global amino acid identity between homologs. (**C**) *aux4* clusters were identified at two locations across the chromosome and megaplasmid, and these locations are shown relative to the GMI1000 replicons. Panel A indicates which genomes encode the cluster at each location. The figure was generated with a combination of KBase BLASTp, iToL, Clinker, and Affinity Designer.

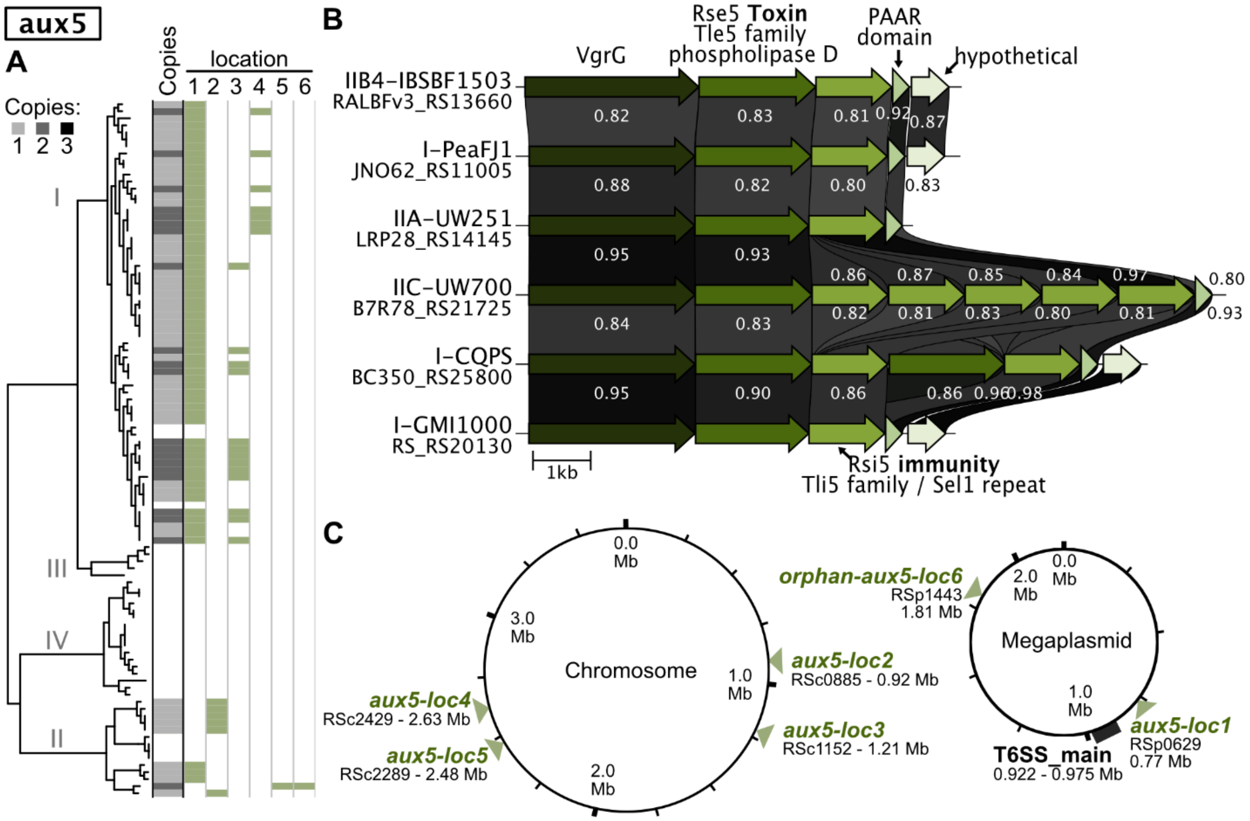


**Figure S8. Phylogenetic distribution, genetic organization/synteny, and chromosomal location of auxiliary *vgrG*-linked cluster 5 (*aux5*).** (**A**) Phylogenetic distribution and copy number of *aux5* across high-quality RSSC genomes. (**B**) Genetic organization/synteny of the cluster from 6 genomes: phyl. I PeaJF1, phyl. I CQPS, phyl. I GMI1000, phyl. IIA UW251, phyl. IIB-4 IBSBF1503, and IIC-7 UW700. *aux5* encodes a VgrG, a Tle5-family phospholipase D, one-or-more Tli5 immunity protein(s) with Sel1 repeats, a PAAR domain protein, and a variably present hypothetical protein. Greyscale links indicate the global amino acid identity between homologs. (**C**) *aux5* clusters were identified at six locations across the chromosome and megaplasmid, and these locations are shown relative to the GMI1000 replicons. Panel A indicates which genomes encode the cluster at each location. The figure was generated with a combination of KBase BLASTp, iToL, Clinker, and Affinity Designer.

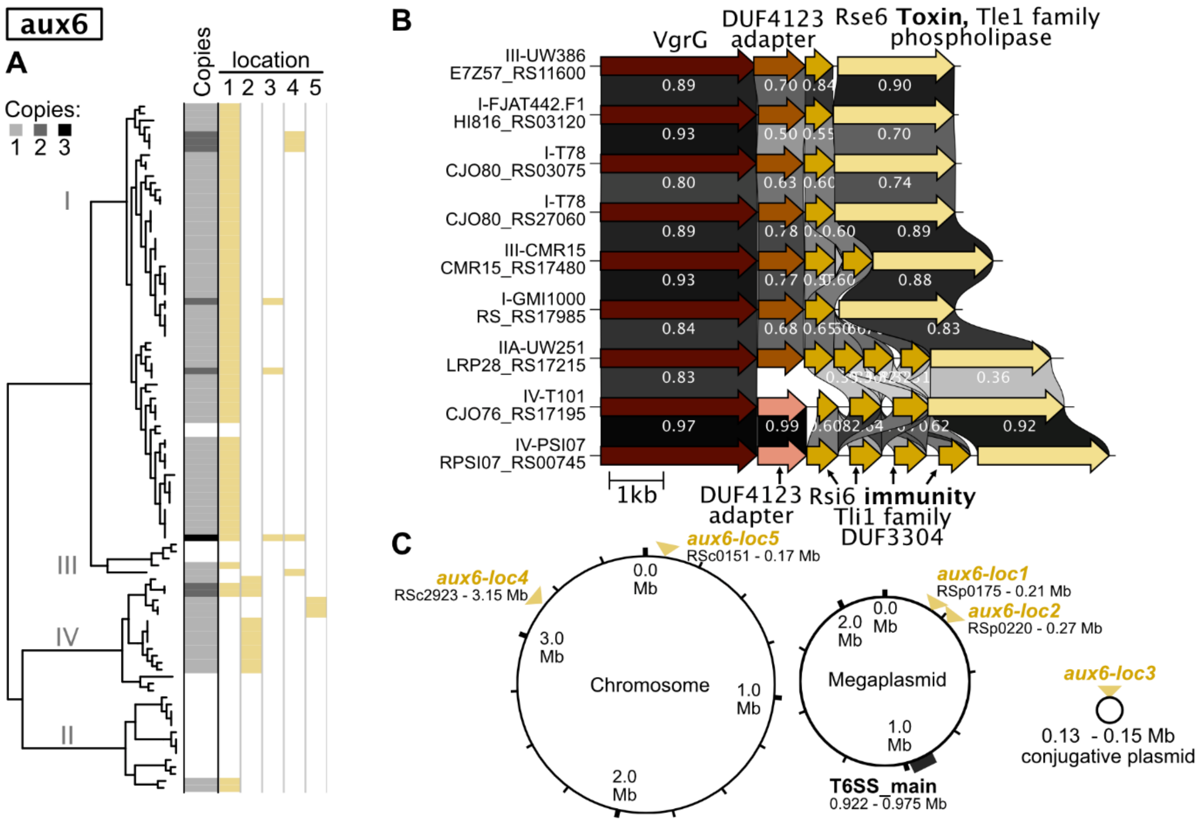


**Figure S9. Phylogenetic distribution, genetic organization/synteny, and chromosomal location of auxiliary *vgrG*-linked cluster 6 (*aux6*).** (**A**) Phylogenetic distribution and copy number of *aux6* across high-quality RSSC genomes. (**B**) Genetic organization/synteny of the cluster from 9 genomes: phyl. I FJAT442.F1, two paralogous clusters from phyl. I T78, phyl. I GMI1000, phyl. IIA UW251, phyl. III CMR15, phyl. III UW386, phyl. IV T101, and phyl. IV PSI07. *aux6* encodes a VgrG, a DUF4123 adaptor protein, one-or-more Tli1 family immunity proteins with a DUF3304 domain, and a Tle1 family phospholipase effector. Greyscale links indicate the global amino acid identity between homologs. (**C**) *aux6* clusters were identified at five locations across the chromosome and megaplasmid, and these locations are shown relative to the GMI1000 replicons. Panel A indicates which genomes encode the cluster at each location. The figure was generated with a combination of KBase BLASTp, iToL, Clinker, and Affinity Designer.

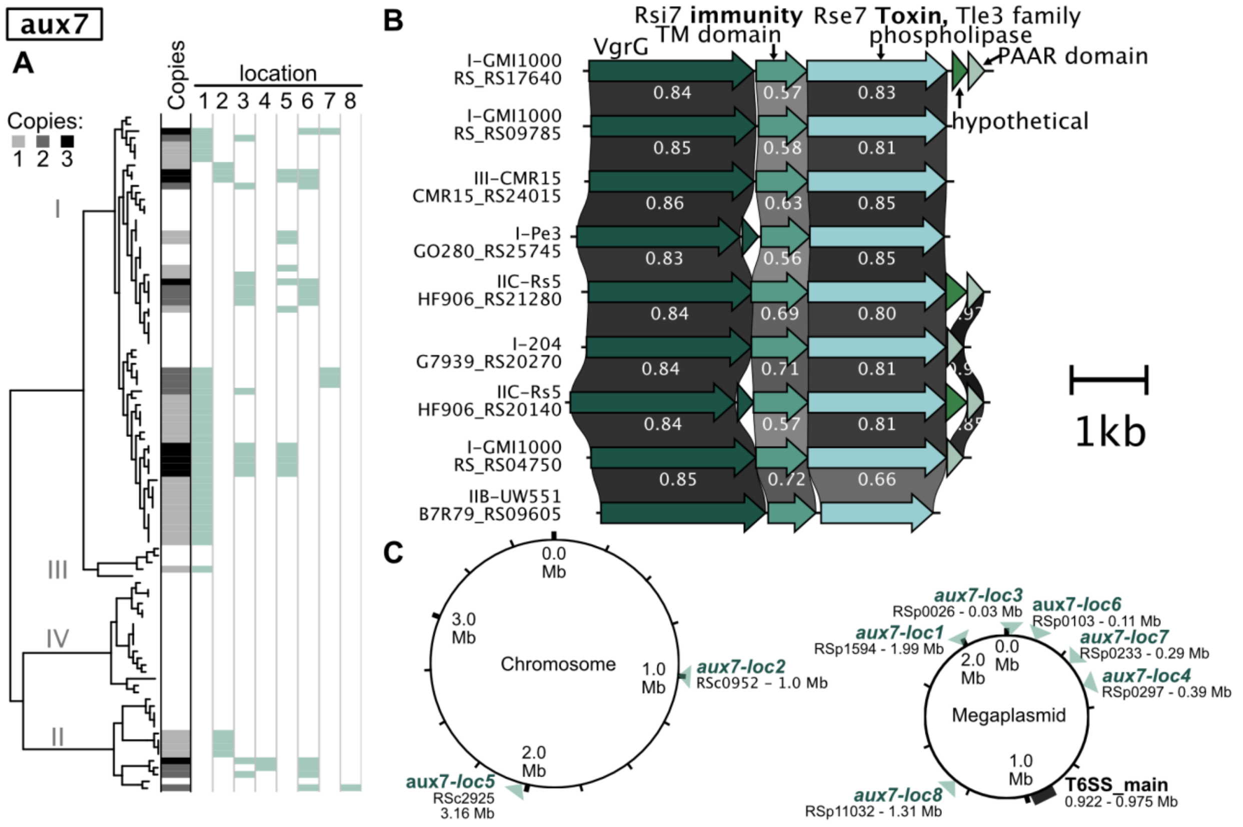


**Figure S10. Phylogenetic distribution, genetic organization/synteny, and chromosomal location of auxiliary *vgrG*-linked cluster 7 (*aux7*).** (**A**) Phylogenetic distribution and copy number of *aux7* across high-quality RSSC genomes. (**B**) Genetic organization/synteny of the cluster from 9 genomes: three distinct paralogous clusters from phyl. I GMI1000, phyl. I Pe3, phyl. I 204, phyl. IIB-1 UW551, two paralogous clusters from phyl. IIC-7 Rs5, and phyl. III CMR15. *aux7* encodes a VgrG, an immunity protein with a transmembrane (TM) domain, a Tle3-family phospholipase effector, a variably present PAAR domain protein, and a variably present hypothetical protein. Greyscale links indicate the global amino acid identity between homologs. (**C**) *aux7* clusters were identified at seven locations across the chromosome and megaplasmid, and these locations are shown relative to the GMI1000 replicons. Panel A indicates which genomes encode the cluster at each location. The figure was generated with a combination of KBase BLASTp, iToL, Clinker, and Affinity Designer.

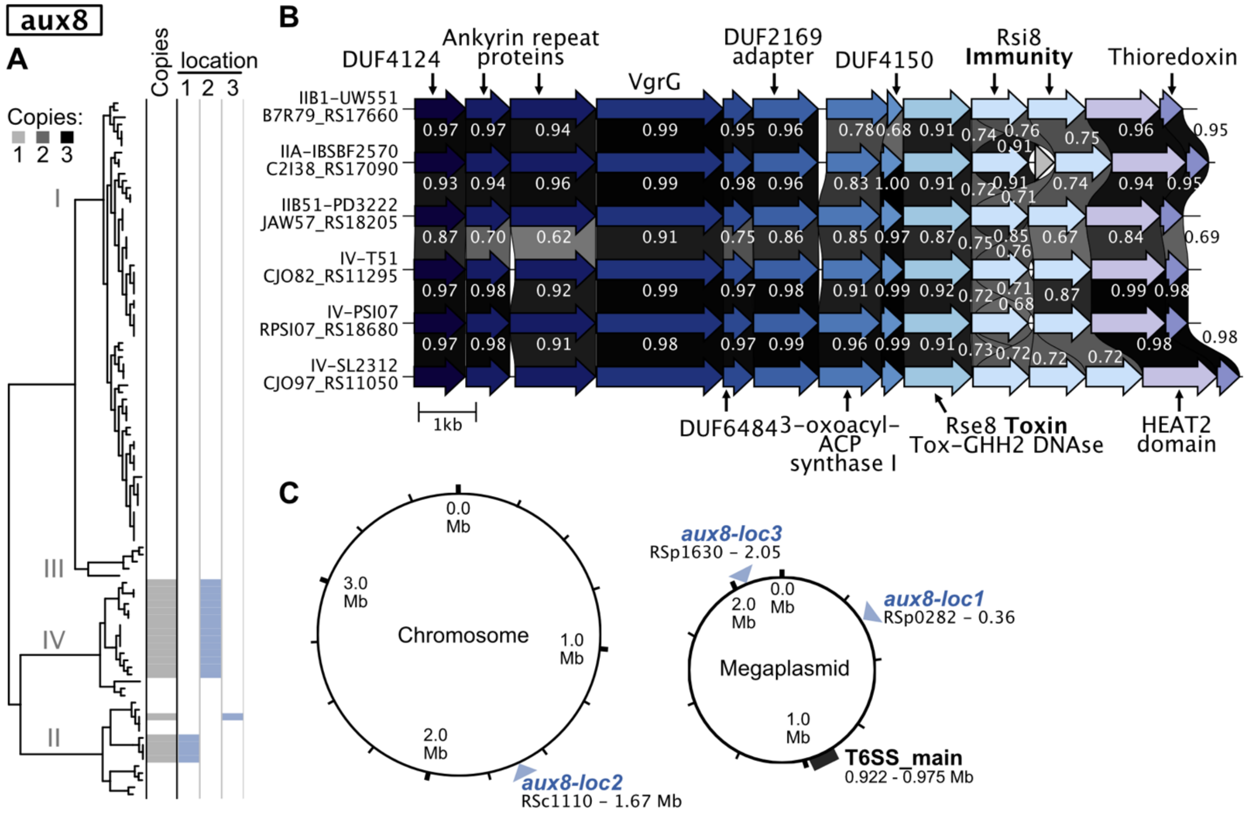


**Figure S11. Phylogenetic distribution, genetic organization/synteny, and chromosomal location of auxiliary *vgrG*-linked cluster 8 (*aux8*).** (**A**) Phylogenetic distribution and copy number of *aux8* across high-quality RSSC genomes. (**B**) Genetic organization/synteny of the cluster from 6 genomes: phyl. IIA IBSBF2570, phyl. IIB-1 UW551, phyl. IIB-51 PD3222, phyl. IV T51, phyl. IV PSI07, and phyl. IV SL2312. *aux8* encodes a DUF4124 protein, two ankyrin repeat proteins, a VgrG, a DUF6484 protein, a DUF2169 adaptor, a 3-oxoacyl-ACP synthase I, a DUF4150 protein, an effector with a Tox-GHH2 DNAse domain, two-or-more immunity proteins, a HEAT2 domain protein, and a thioredoxin. Greyscale links indicate the global amino acid identity between homologs. (**C**) *aux8* clusters were identified at three locations across the chromosome and megaplasmid, and these locations are shown relative to the GMI1000 replicons. Panel A indicates which genomes encode the cluster at each location. The figure was generated with a combination of KBase BLASTp, iToL, Clinker, and Affinity Designer.

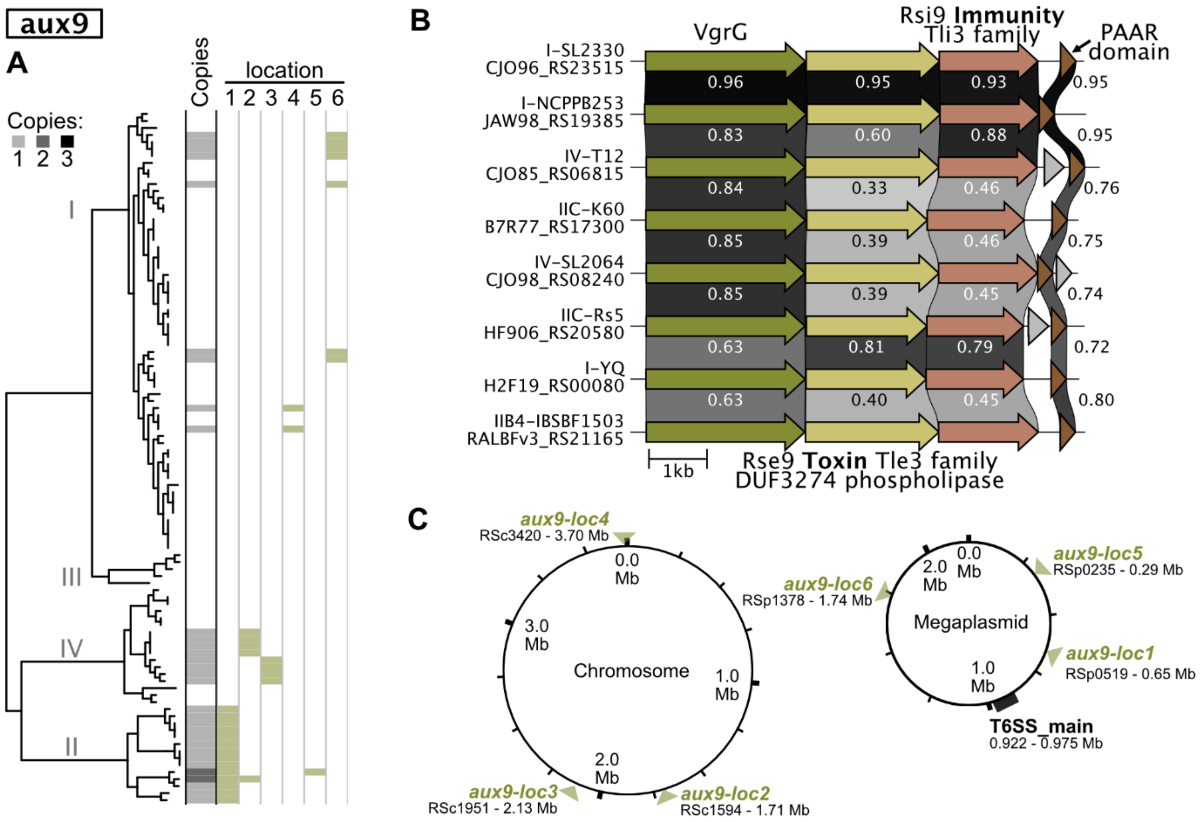


**Figure S12. Phylogenetic distribution, genetic organization/synteny, and chromosomal location of auxiliary *vgrG*-linked cluster 9 (*aux9*).** (**A**) Phylogenetic distribution and copy number of *aux9* across high-quality RSSC genomes. (**B**) Genetic organization/synteny of the cluster from eight genomes: phyl. I SL2330, phyl. I NCPPB253, phyl. I YQ, phyl IIB-4 IBSBF1503, phyl. IIC-7 K60, phyl. IIC-7 Rs5, and phyl. IV SL2064. *aux9* encodes a VgrG, a Tle3-family phospholipase with a DUF3274 domain, a Tli3 immunity protein(s), a PAAR domain protein, and variably present hypothetical proteins. Greyscale links indicate the global amino acid identity between homologs. (**C**) *aux9* clusters were identified at six locations across the chromosome and megaplasmid, and these locations are shown relative to the GMI1000 replicons. Panel A indicates which genomes encode the cluster at each location. The figure was generated with a combination of KBase BLASTp, iToL, Clinker, and Affinity Designer.

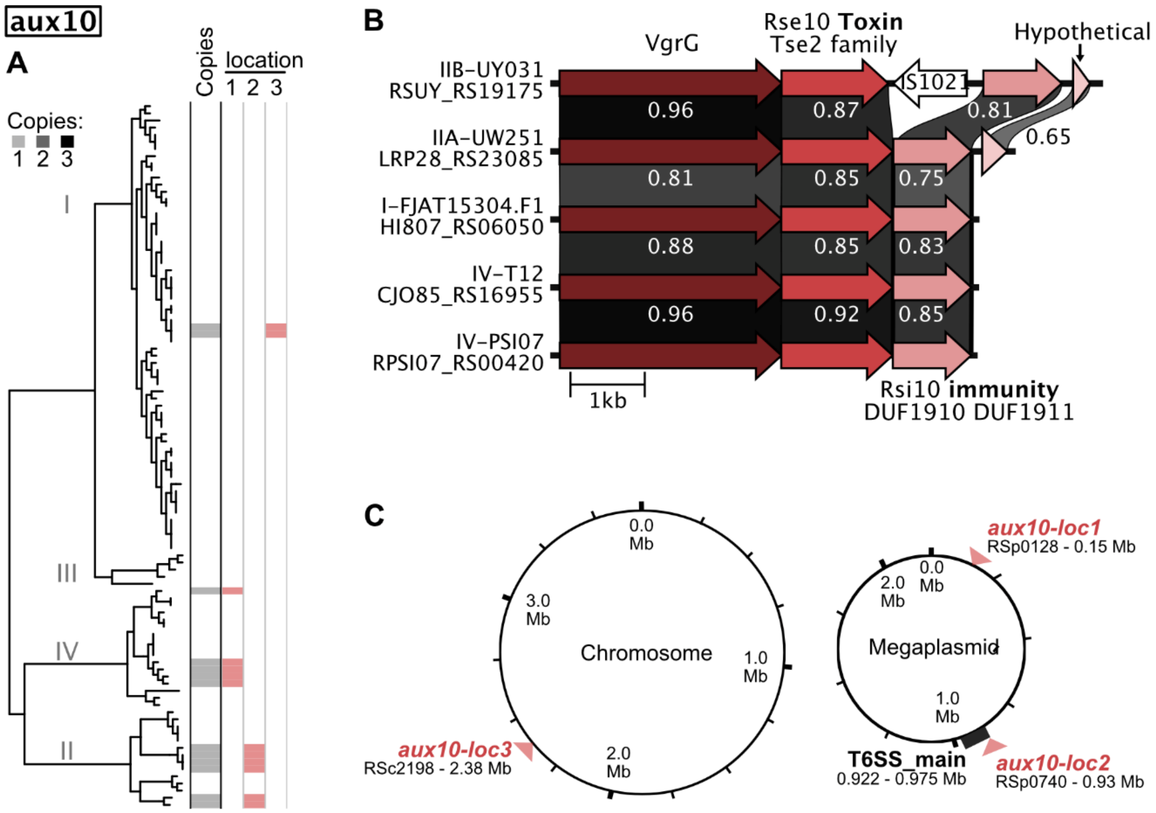


**Figure S13. Phylogenetic distribution, genetic organization/synteny, and chromosomal location of auxiliary *vgrG*-linked cluster 10 (*aux10*).** (**A**) Phylogenetic distribution and copy number of *aux10* across high-quality RSSC genomes. (**B**) Genetic organization/synteny of the cluster from 5 genomes: phyl. I FJAT15304.F1, phyl. IIA UW251, phyl. IIB-1 UY031, phyl. IV T12, and phyl. IV-10 PSI07. *aux10* encodes a VgrG, a Tse2-family effector with an unknown target, an immunity protein with DUF1910 and DUF1911 domains, and a variably present hypothetical protein. Greyscale links indicate the global amino acid identity between homologs. (**C**) *aux10* clusters were identified at three locations across the chromosome and megaplasmid, and these locations are shown relative to the GMI1000 replicons. Panel A indicates which genomes encode the cluster at each location. The figure was generated with a combination of KBase BLASTp, iToL, Clinker, and Affinity Designer.

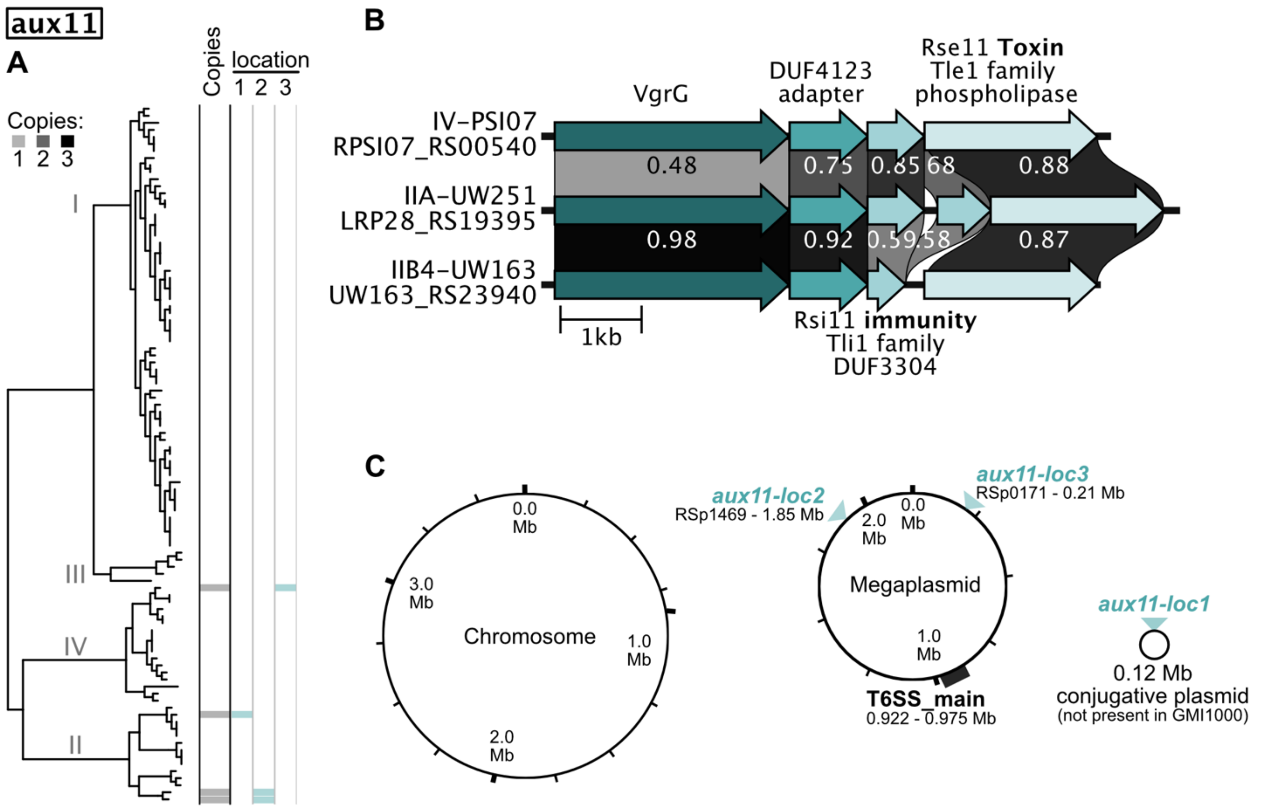


**Figure S14. Phylogenetic distribution, genetic organization/synteny, and chromosomal location of auxiliary *vgrG*-linked cluster 11 (*aux11*).** (**A**) Phylogenetic distribution and copy number of *aux11* across high-quality RSSC genomes. (**B**) Genetic organization/synteny of the cluster from 3 genomes: phyl. IIA UW251, phyl. IIB-4 UW163, and IV PSI07. *aux11* encodes a VgrG, a DUF4123 adapter, one-or-more Tli1 family immunity proteins with a DUF3304 domain, and a Tle1 family phospholipase effector. Greyscale links indicate the global amino acid identity between homologs. (**C**) *aux11* clusters were identified at three locations across the chromosome, megaplasmid, and conjugative accessory plasmids, and these locations are shown relative to the GMI1000 replicons. Panel A indicates which genomes encode the cluster at each location. The figure was generated with a combination of KBase BLASTp, iToL, Clinker, and Affinity Designer.

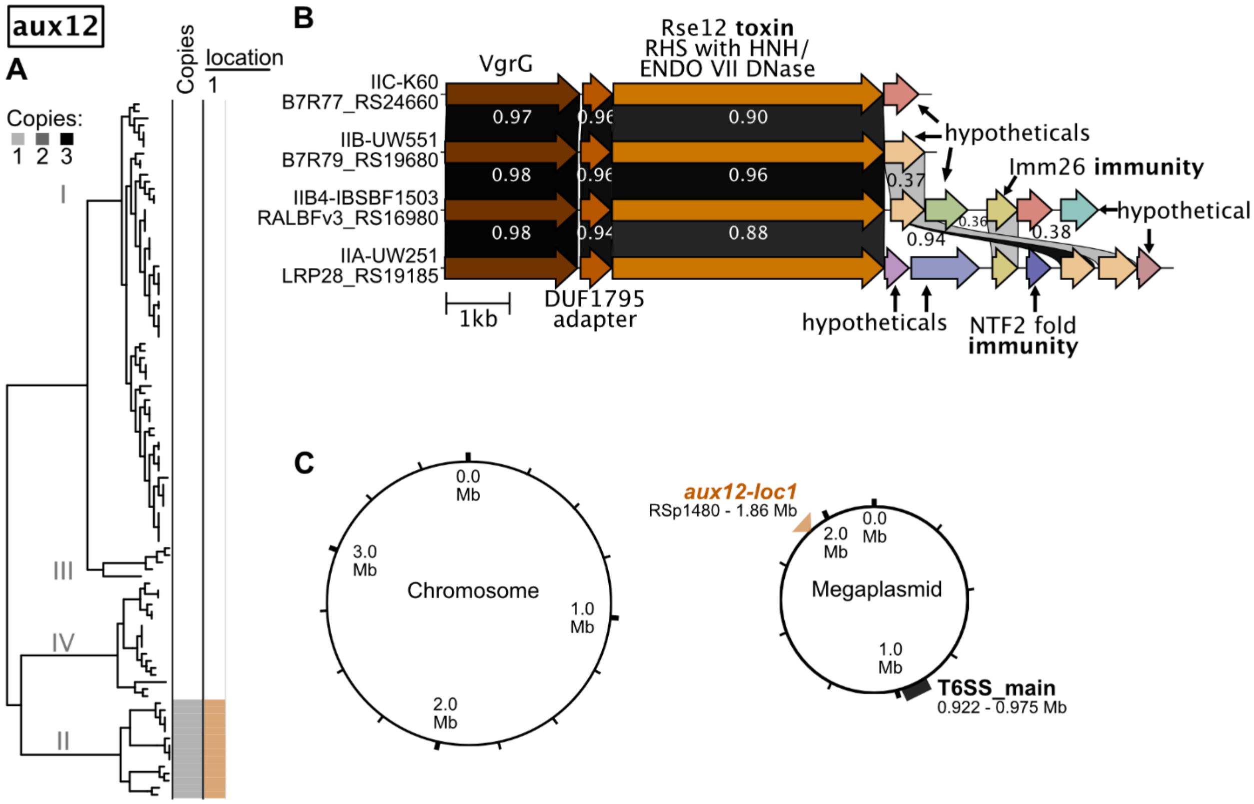


**Figure S15. Phylogenetic distribution, genetic organization/synteny, and chromosomal location of auxiliary *vgrG*-linked cluster 12 (*aux12*).** (**A**) Phylogenetic distribution and copy number of *aux12* across high-quality RSSC genomes. (**B**) Genetic organization/synteny of the cluster from 4 genomes: phyl. IIA UW251, phyl. IIB-1 UW551, phyl. IIB-4 IBSBF1503, and phyl. IIC-7 K60. *aux12* encodes a VgrG, a DUF1795 adapter, an RHS protein with a C-terminal HNH/ENDO VII DNase effector, and an array of one-or-more variably present proteins encoding hypotheticals, Imm26 family immunity proteins, and an NFT2-fold immunity protein. The small genes downstream of the RHS effector likely encode an array of immunity proteins. Greyscale links indicate the global amino acid identity between homologs. (**C**) *aux12* clusters were identified at one location of the megaplasmid. The location is shown relative to the GMI1000 megaplasmid. The figure was generated with a combination of KBase BLASTp, iToL, Clinker, and Affinity Designer.

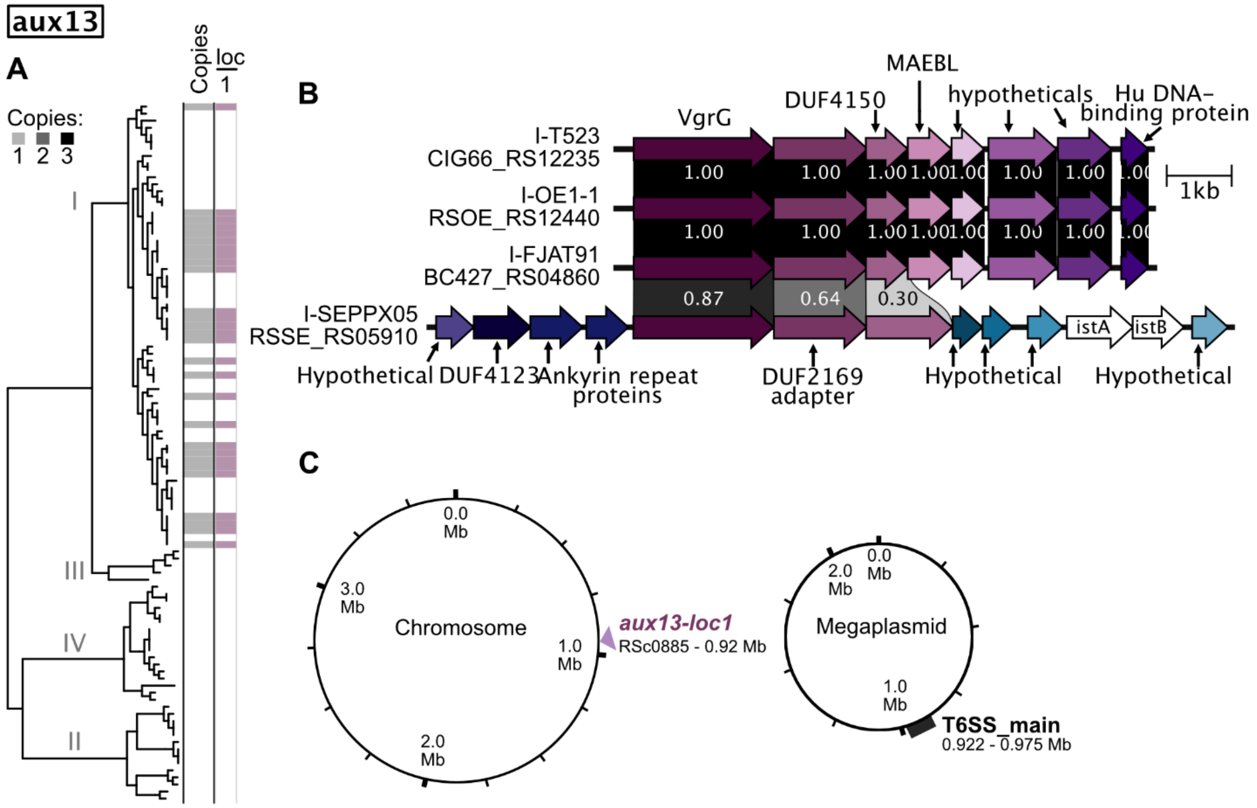


**Figure S16. Phylogenetic distribution, genetic organization/synteny, and chromosomal location of auxiliary *vgrG*-linked cluster 13 (*aux13*).** (**A**) Phylogenetic distribution and copy number of *aux13* across high-quality RSSC genomes. (**B**) Genetic organization/synteny of the cluster from 4 genomes: phyl. I T523, phyl. I OE1-1, phyl. I-FJAT91, and phyl. I SEPPX05. *aux13* encodes a VgrG, a DUF2169 family adapter, a DUF4150 protein, four hypothetical proteins, and a Hu DNA-binding protein. Greyscale links indicate the global amino acid identity between homologs. (**C**) The *aux13* clusters were identified at one location on the chromosome, and these locations are shown relative to the GMI1000 replicons. Panel A indicates which genomes encode the cluster at each location. The figure was generated with a combination of KBase, BLASTp, iToL, Clinker, and Affinity Designer.

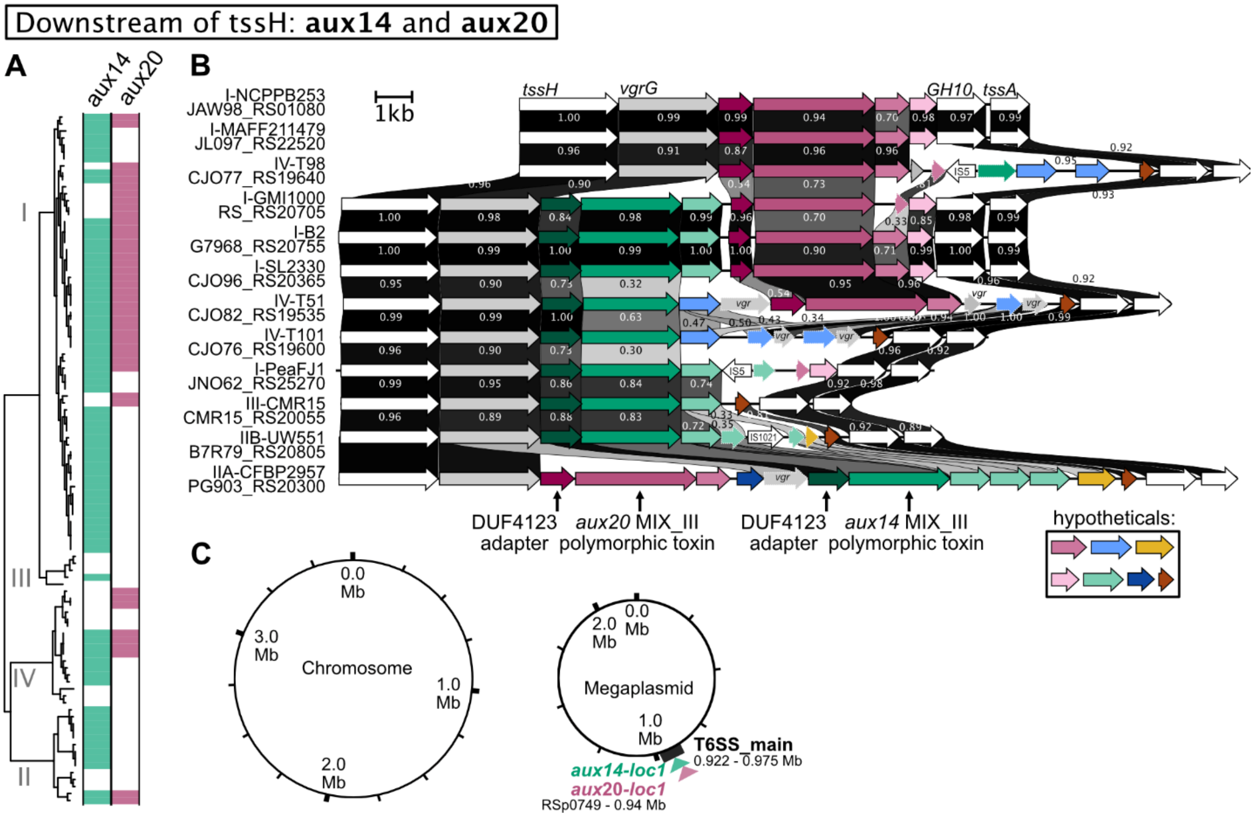


**Figure S17. Phylogenetic distribution, genetic organization/synteny, and chromosomal location of auxiliary *vgrG*-linked clusters located between *tssH* and GH10/*tssA* (*aux14* and *aux20*).** (**A**) Phylogenetic distribution of *aux14* and *aux20* across high-quality RSSC genomes. (**B**) Genetic organization/synteny of four clusters that only contain *aux14*, three clusters that only contain *aux20*, four that contain *aux14* upstream of *aux20*, and one that contains *aux20* upstream of *aux14*. The clusters are from phyl. I NCPPB253, phyl. I MAFF211479, phyl. I GMI1000, phyl. I B2, phyl. I SL2330, phyl. I peaFJ1, phyl. IIA CDBP2957, phyl. IIB1 UW551, phyl. III CMR15, phyl. IV T98, phyl. IV T51, and phyl. IV T101. *aux14* encodes a VgrG, a DUF4123 adaptor, a MIX_III polymorphic effector with unknown C-terminal effector, and a hypothetical protein (presumed immunity). *Aux20* encodes a VgrG, a DUF4123 adaptor, a MIX_III polymorphic effector with unknown C-terminal effector, and a hypothetical protein (presumed immunity). (**C**) These clusters are only encoded at the T6SS main locus on the megaplasmid, and this location is shown relative to the GMI1000 replicons. The figure was generated with a combination of KBase BLASTp, iToL, Clinker, and Affinity Designer.

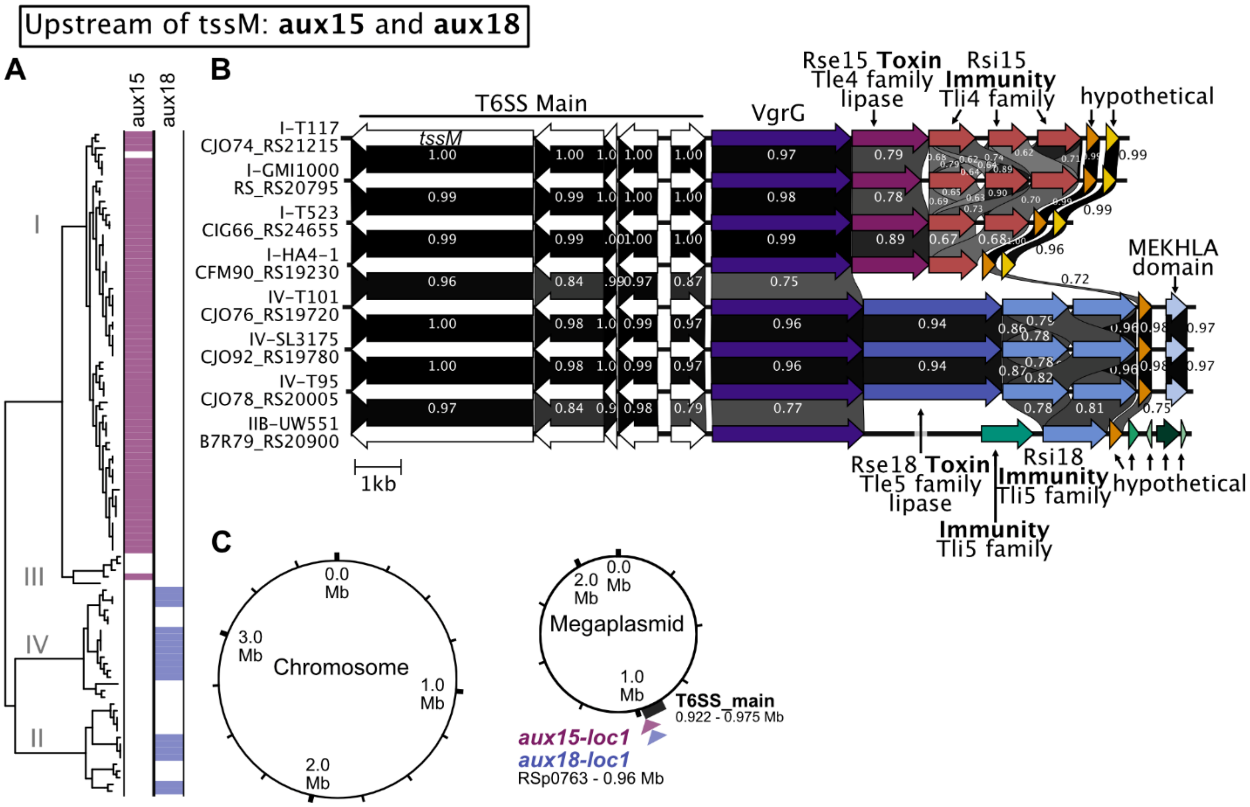


**Figure S18. Phylogenetic distribution, genetic organization/synteny, and chromosomal location of auxiliary *vgrG*-linked clusters located upstream of *tssM* (*aux15* and *aux18*).** (**A**) Phylogenetic distribution of *aux15* and *aux18* across high-quality RSSC genomes. (**B**) Genetic organization/synteny of four *aux15* clusters, three *aux18* clusters, and clusters lacking effectors. The *aux15* clusters are from phyl. I T117, phyl. I GMI1000, phyl. I T523, and phyl. I HA4-1. The *aux18* clusters are from phyl. IV T101, phyl. IV SL3175, and phyl. IV T95. The cluster without an effector is from phyl. IIB-1 UW551. *aux15* encodes a VgrG, a Tle4 family phospholipase, one-or-more Tli4 family immunity protein(s), and two hypothetical proteins. *aux18* encodes a VgrG, a Tle5 phospholipase effector, two Tli5 immunity proteins, a hypothetical protein that is also encoded in *aux15*, and a MEKHLA domain protein. The IIB-1 cluster encodes two distinct Tli5 immunity proteins, the hypothetical protein from *aux15/aux18* and an array of additional hypothetical proteins. (**C**) These clusters are only encoded at the T6SS main locus on the megaplasmid, and this location is shown relative to the GMI1000 replicons. The figure was generated with a combination of KBase, BLASTp, iToL, Clinker, and Affinity Designer.

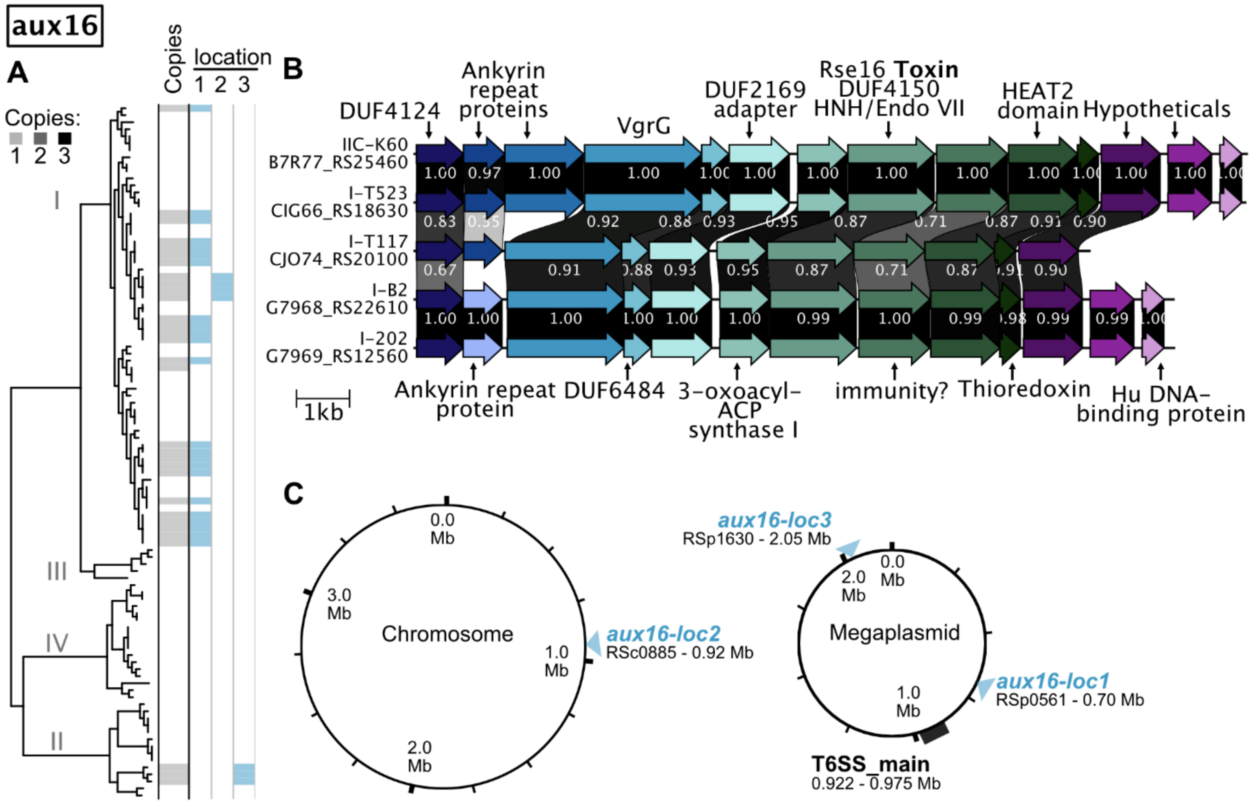


**Figure S19. Phylogenetic distribution, genetic organization/synteny, and chromosomal location of auxiliary *vgrG*-linked cluster 16 (*aux16*).** (**A**) Phylogenetic distribution and copy number of *aux16* across high-quality RSSC genomes. (**B**) Genetic organization/synteny of the cluster from 5 genomes: phyl. I T523, phyl. I T117, phyl. I B2, phyl. I 202 and phyl. IIC-7 K60. *aux16* encodes a DUF4123 protein, one-to-two ankyrin repeat proteins, a VgrG, a DUF6484 protein, a DUF2169 adaptor, a 3-oxoacyl-ACP synthase I, a DUF4150 protein with a Tox-GHH2 DNAse domain, an immunity protein, a HEAT2 domain protein, thioredoxin, one-or-more hypothetical, and a variably present Hu DNA binding protein. Greyscale links indicate the global amino acid identity between homologs. (**C**) *aux16* clusters were identified at three locations across the chromosome and megaplasmid, and these locations are shown relative to the GMI1000 replicons. Panel A indicates which genomes encode the cluster at each location. The figure was generated with a combination of KBase, BLASTp, iToL, Clinker, and Affinity Designer.

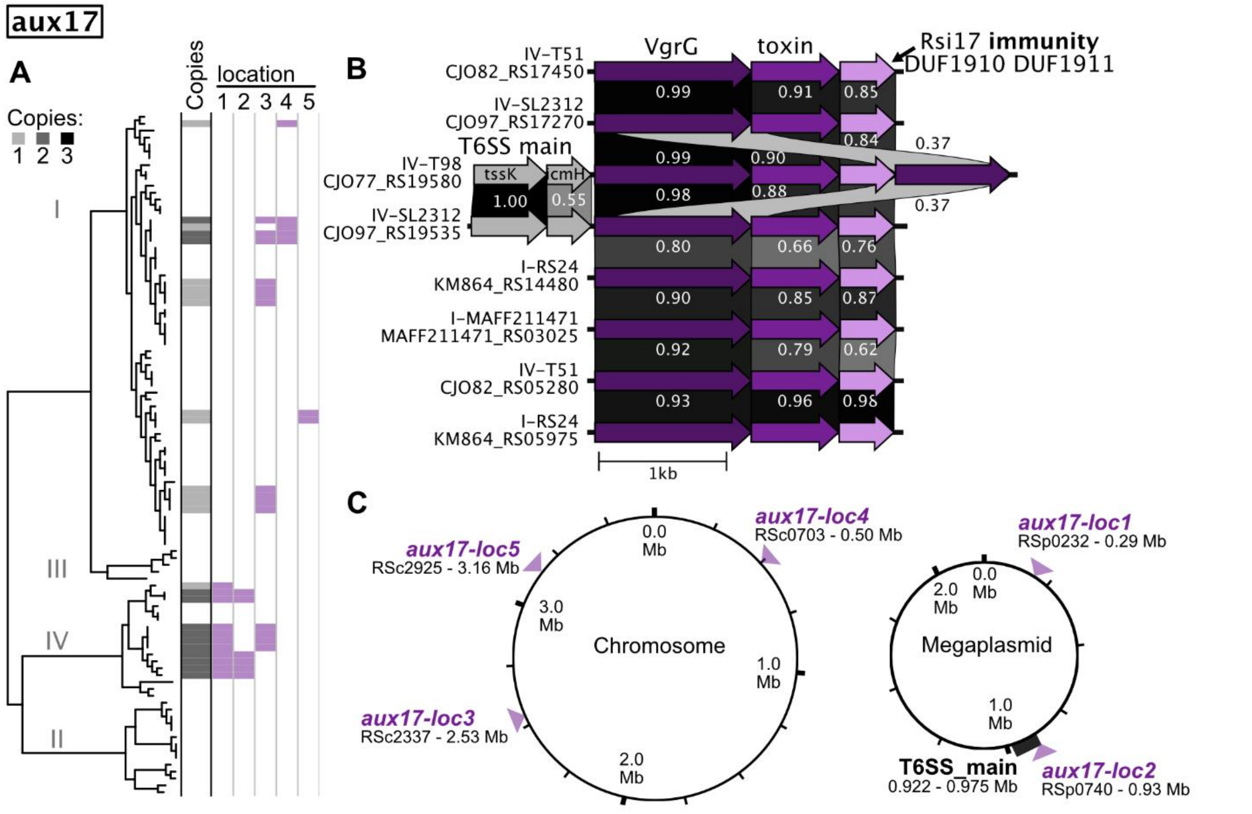


**Figure S20. Phylogenetic distribution, genetic organization/synteny, and chromosomal location of auxiliary *vgrG*-linked cluster 17 (*aux17*).** (**A**) Phylogenetic distribution and copy number of *aux17* across high-quality RSSC genomes. (**B**) Genetic organization/synteny of 8 clusters: phyl. I MAFF211471, two paralogous clusters from phyl. I RS24, two paralogous clusters from phyl. IV T51, two paralogous clusters from phyl. IV SL2312, and phyl. IV T98. *aux17* encodes a VgrG, a putative effector that lacks any identified domains, and an immunity protein with DUF1910 and DUF1911 domains. Greyscale links indicate the global amino acid identity between homologs. (**C**) *aux17* clusters were identified at five locations across the chromosome and megaplasmid, and these locations are shown relative to the GMI1000 replicons. Panel A indicates which genomes encode the cluster at each location. The figure was generated with a combination of KBase, BLASTp, iToL, Clinker, and Affinity Designer.

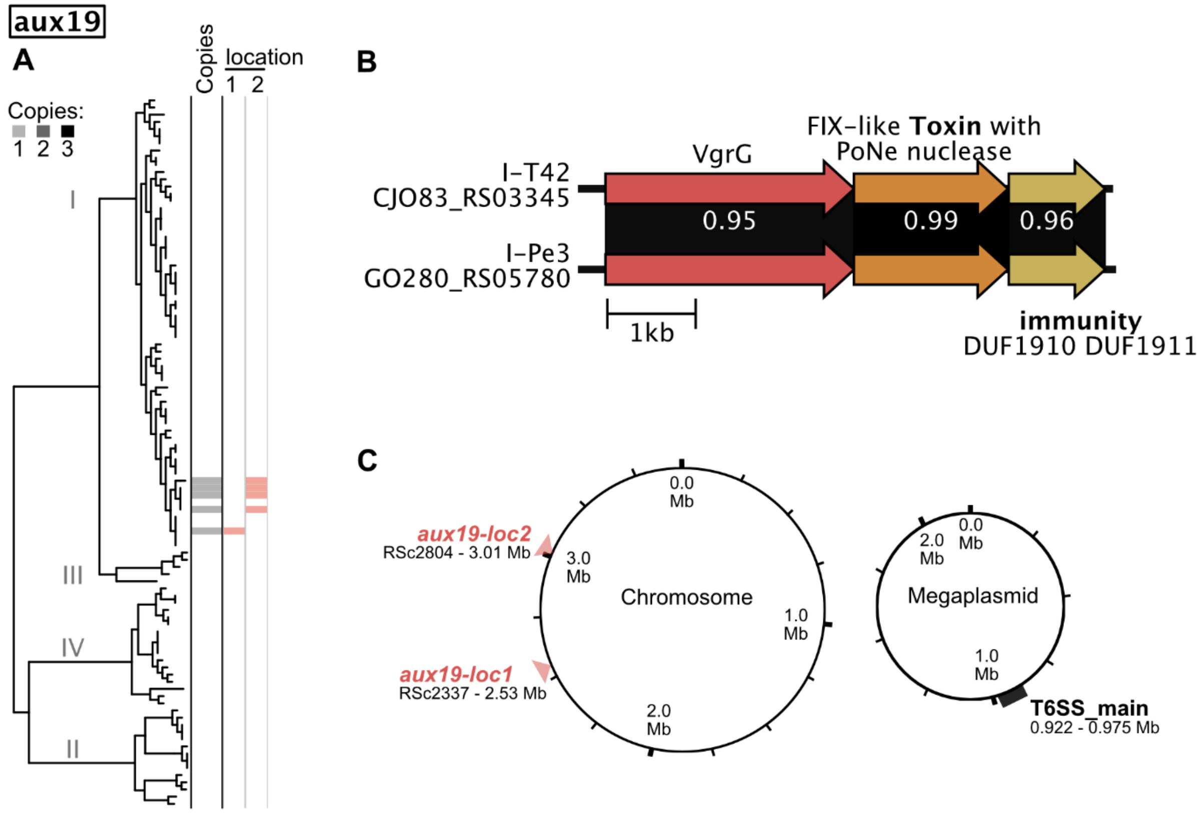


**Figure S21. Phylogenetic distribution, genetic organization/synteny, and chromosomal location of auxiliary *vgrG*-linked cluster 19 (*aux19*).** (**A**) Phylogenetic distribution and copy number of *aux19* across high-quality RSSC genomes. (**B**) Genetic organization/synteny of the cluster from two genomes: phyl I T42, and phyl. I Pe3. *aux19* encodes a VgrG, a FIX-like polymorphic effector with a C-terminal PoNe nuclease domain, and an immunity protein with DUF1910 and DUF1911 domains. Greyscale links indicate the global amino acid identity between homologs. (**C**) *aux19* clusters were identified at two locations on the chromosome, and these locations are shown relative to the GMI1000 replicon. Panel A indicates which genomes encode the cluster at each location. The figure was generated with a combination of KBase, BLASTp, iToL, Clinker, and Affinity Designer.

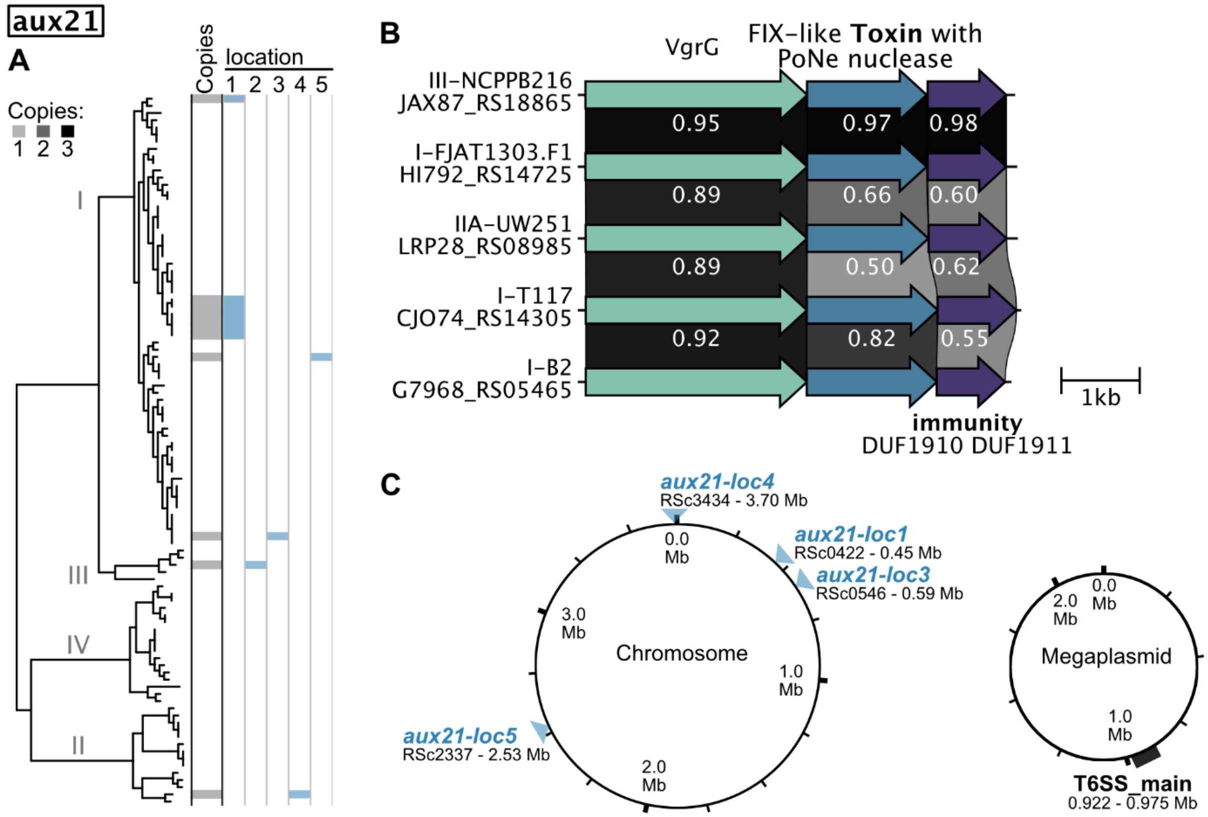


**Figure S22. Phylogenetic distribution, genetic organization/synteny, and chromosomal location of auxiliary *vgrG*-linked cluster 21 (*aux21*).** (**A**) Phylogenetic distribution and copy number of *aux21* across high-quality RSSC genomes. (**B**) Genetic organization/synteny of the cluster from five genomes: phyl. I T117, phyl. I B2, phyl. I FJAT1303.F1, phyl. IIA UW251, and phyl. III NCPPB216. *aux21* encodes a VgrG, a FIX-like polymorphic effector with a SEN1 helicase domain and a C-terminal PoNe nuclease domain, and a DUF1910/DUF1911 immunity protein. Greyscale links indicate the global amino acid identity between homologs. (**C**) *aux21* clusters were identified at five locations across the chromosome, and these locations are shown relative to the GMI1000 replicon. Panel A indicates which genomes encode the cluster at each location. The figure was generated with a combination of KBase, BLASTp, iToL, Clinker, and Affinity Designer.

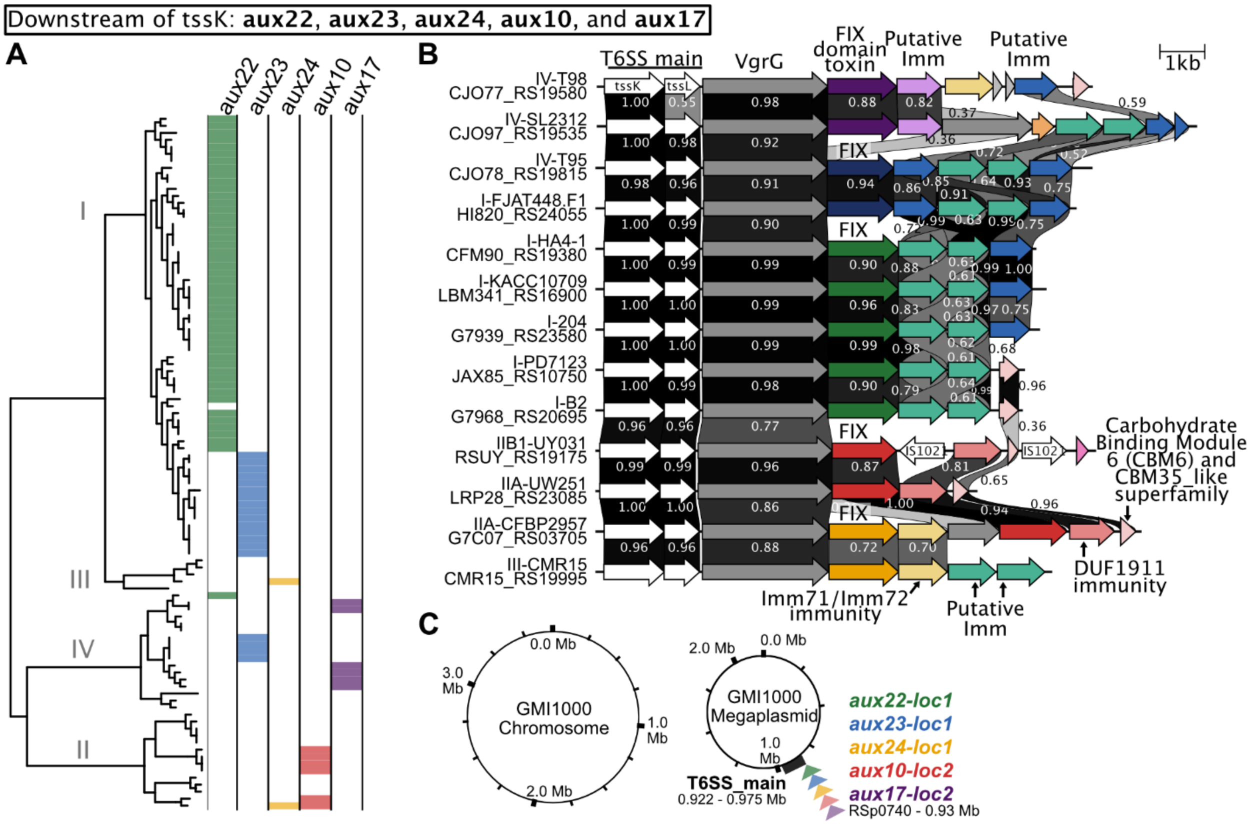


**Figure S23. Phylogenetic distribution, genetic organization/synteny, and chromosomal location of auxiliary *vgrG*-linked clusters that are downstream of *tssJKL* (*aux22, aux 23, aux24, aux10,* and *aux17*).** (**A**) Phylogenetic distribution of each cluster across high-quality RSSC genomes. (**B**) Genetic organization/synteny of 13 clusters. All clusters encode a VgrG, one of five distinct FIX domain effectors, and one or more putative immunity proteins. *aux10* is represented by phyl. IIB-1 UY031 and phyl. IIA UW251. *aux17* is represented by phyl. IV T98 and phyl. IV SL2312. *aux22* is represented by phyl. I strains HA4-1, 204, PD7123, and B2. *aux23* is represented by phyl. IV T95 and phyl. I FJAT448.F1. *aux24* is represented by phyl. IIA CFBP2957 and phyl. III CMR15. Greyscale links indicate the global amino acid identity between homologs. (**C**) Three of these clusters (*aux22*, *aux23*, and *aux24*) are encoded exclusively at the T6SS main locus on the megaplasmid. *aux10* and *aux17* can be found at the T6SS main locus or at the other location indicated on figures S13 and S27 that focus on each of these clusters. The figure was generated with a combination of KBase, BLASTp, iToL, Clinker, and Affinity Designer.

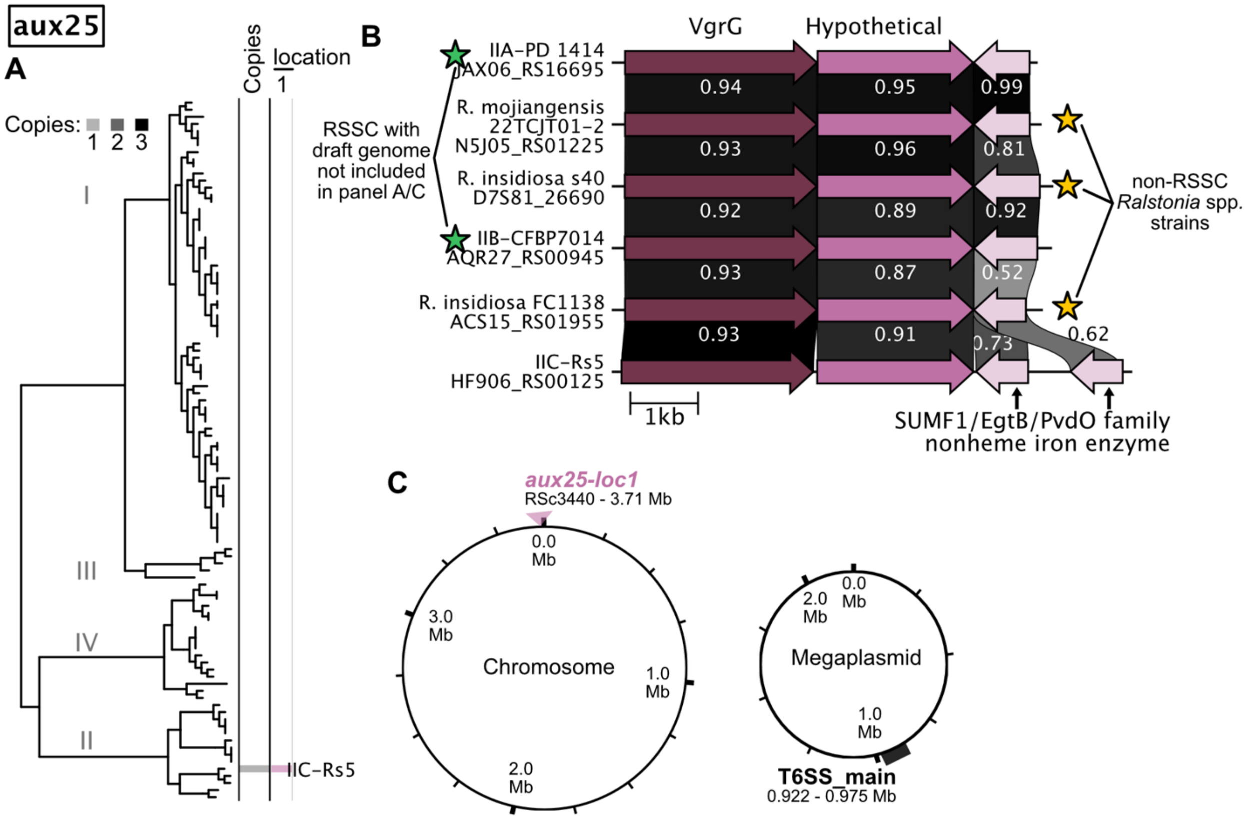


**Figure S24. Phylogenetic distribution, genetic organization/synteny, and chromosomal location of the atypical, auxiliary *vgrG*-linked gene cluster *aux25*.** (**A**) Phylogenetic distribution of *aux25* across high-quality RSSC genomes. Not shown: *aux25* is also encoded in the draft genomes of IIA strain PD1414 and IIB-51 strain CFBP7014. (**B**) Genetic organization/synteny of *aux25* from RSSC strains (IIC Rs5, IIA PD1414, and IIB-51 CFBP7014) and strains in other *Ralstonia* species (FC1138, 22TCJT01-2, and s40). All clusters encode a VgrG, a hypothetical protein, and a SUMF1/EgtB/PvdO family nonheme iron enzyme. The SUMF1 gene is encoded on the opposing strand of the rest of the cluster, which is atypical. Greyscale links indicate the global amino acid identity between homologs. (**C**) The relative location of *aux25 in Rs5* is shown relative to the GMI1000 chromosome. The figure was generated with a combination of KBase, BLASTp, iToL, Clinker, and Affinity Designer.

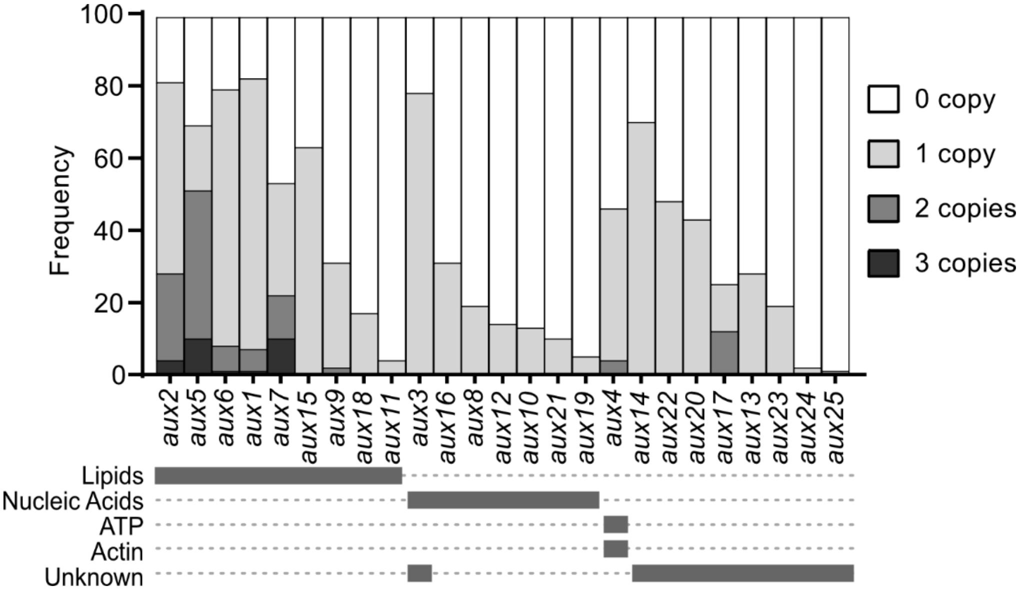


**Fig S25. RSSC genomes encode 0-3 copies of *aux* clusters.** *aux* clusters are organized by the substrate that the toxin targets.

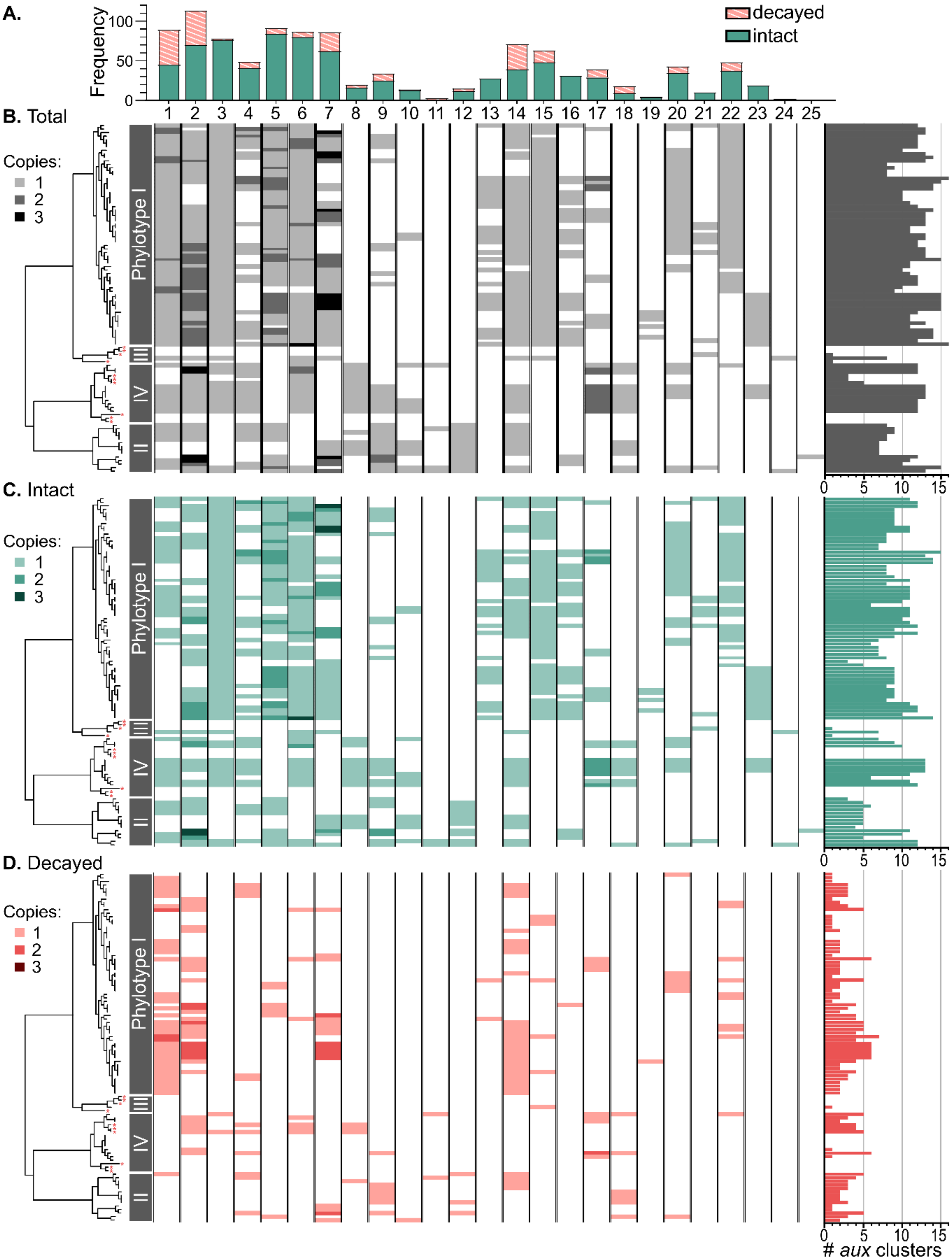


**Figure S26. RSSC strains vary in their repertoires of *vgr*-linked auxiliary toxin/immunity clusters.** The presence of each cluster (*aux1-aux25*) was determined across complete or nearly complete RSSC genomes by a combination of BlastP and Clinker analyses and visualized with iToL. (**A**) shows the frequency of each cluster and the proportion that are intact (genes lack any obvious loss-of-function mutations) or decayed (one-or-more genes have loss-of-function mutations like pseudogenization/frameshifts, deletions that truncate or remove a gene, or transposon insertions into a gene). (**B**) shows the total prevalence of aux clusters in each genome, (**C**) shows the prevalence of intact clusters, and (**D**) shows the prevalence of decayed clusters. The phylogenetic tree was constructed with the KBase "Species Tree" app. Genomes that lack a T6SS are indicated by red asterisks to the right of the branch.

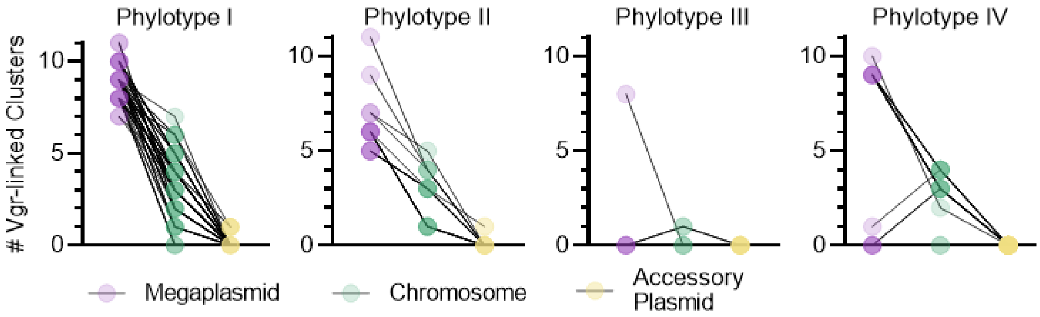


**Figure S27. *aux* clusters are enriched on the megaplasmid for all T6SS⁺ RSSC strains.** Each genome is represented by a line that connects semi-transparent circles. *aux* clusters are more enriched on the megaplasmid than on the chromosome and accessory plasmid of all four RSSC phylotypes. *aux* clusters are dominantly found in Phyl. I and Phyl. II strains in contrast to Phyl. III and Phyl. IV. This could be due to the low sampling of Phyl. III and Phyl. IV strains, thus sequencing more strains from these phylotypes could reveal more information about *aux* clusters content and distribution.

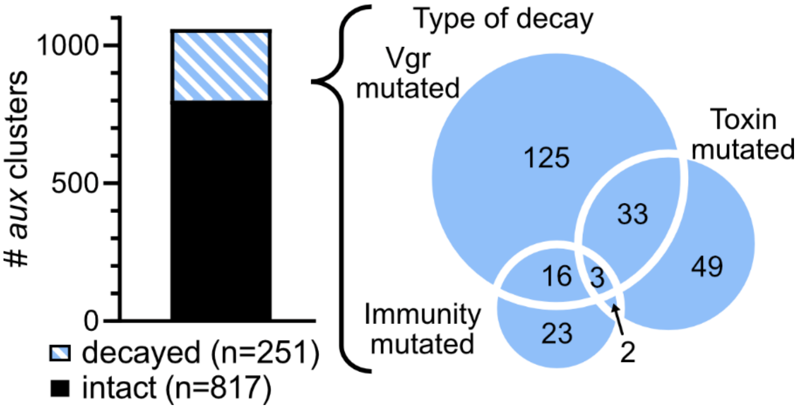


**Figure S28. Patterns of loss-of-function mutations observed in *aux* clusters.** Evidence of gene decay / loss was investigated for each of the 1069 *aux* clusters. Frameshift and other pseudogenization mutations were identified based on NCBI RefSeq annotations. Genetic fragmentation, disruption by transposon insertion, and gene loss were identified through Clinker analysis of synteny. BioVenn (111) was used to create a proportional Venn diagram to reflect whether mutations occurred in the *vgr,* toxin, or immunity gene.

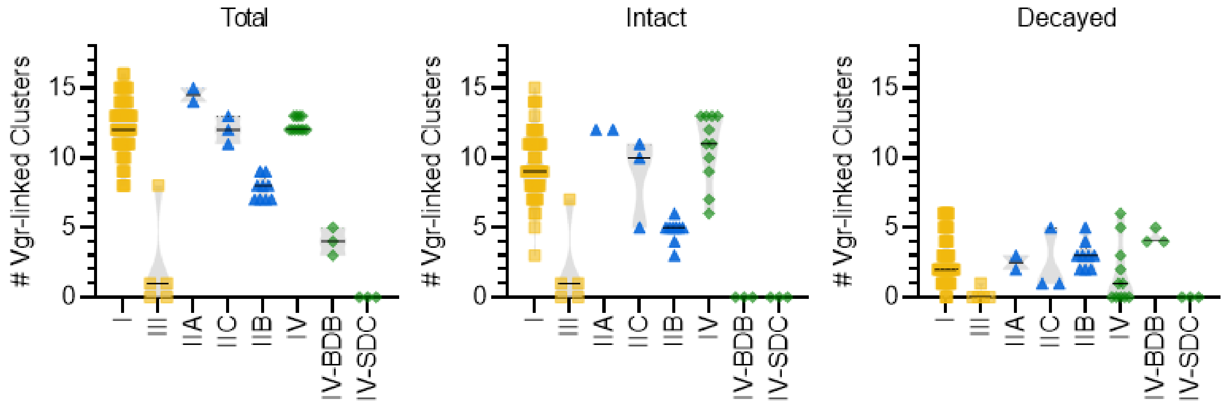


**Figure S29. Comparison of total, intact, and decayed *aux* clusters amongst RSSC clades.**

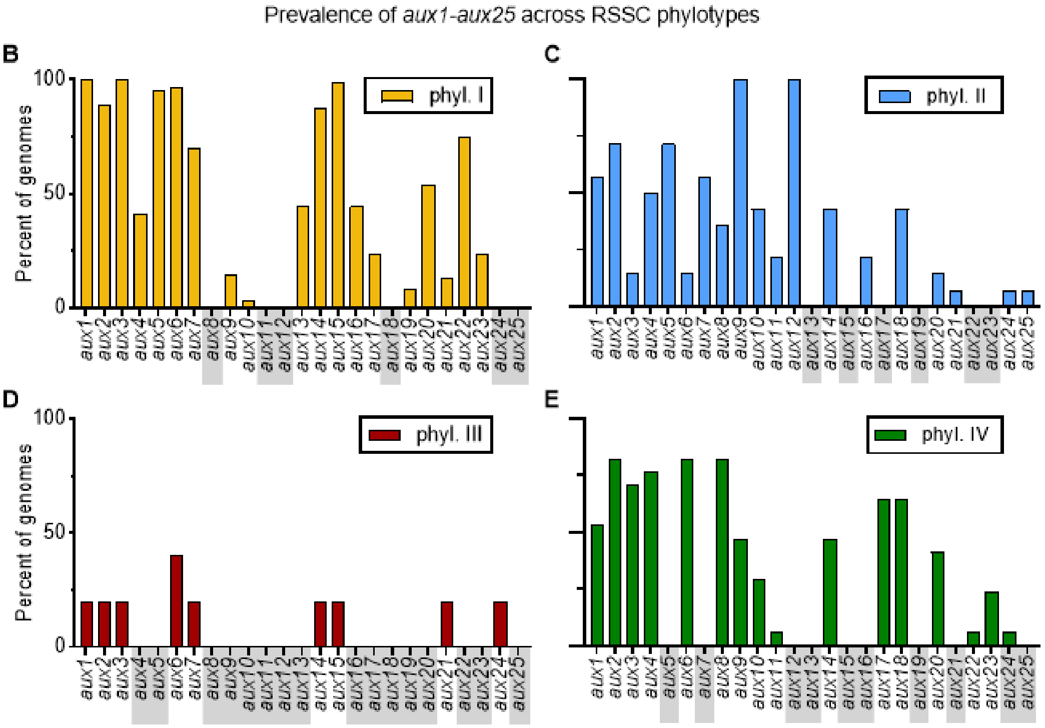


**Figure S30. RSSC lineages vary in their repertoires of auxiliary vgr-linked toxin/antitoxin clusters (*aux* clusters).** *aux* clusters were identified in complete or nearly complete genomes by BLAST searches for genes encoding Vgr, toxins, or immunity proteins. The *aux* clusters were classified by synteny analysis with Clinker. (**A**) RSSC lineages vary in their abundance of *aux* clusters. SDC and BDB refer to the Sumatra Disease of Clove and Banana Blood Disease lineages of phylotype IV, which are transmitted through xylem-sap feeding insect vectors (SDC) or mechanically vectored through tools or pollinating insects (BDB). (**B-E**) *aux* clusters vary in their prevalence in different RSSC phylotypes (n=775 phylotype I, n=135 phyl. II, n=10 phyl. III, n=149 phyl IV). Grey rectangles identify *aux* clusters that were not identified in the genomes of strains in each phylotype.

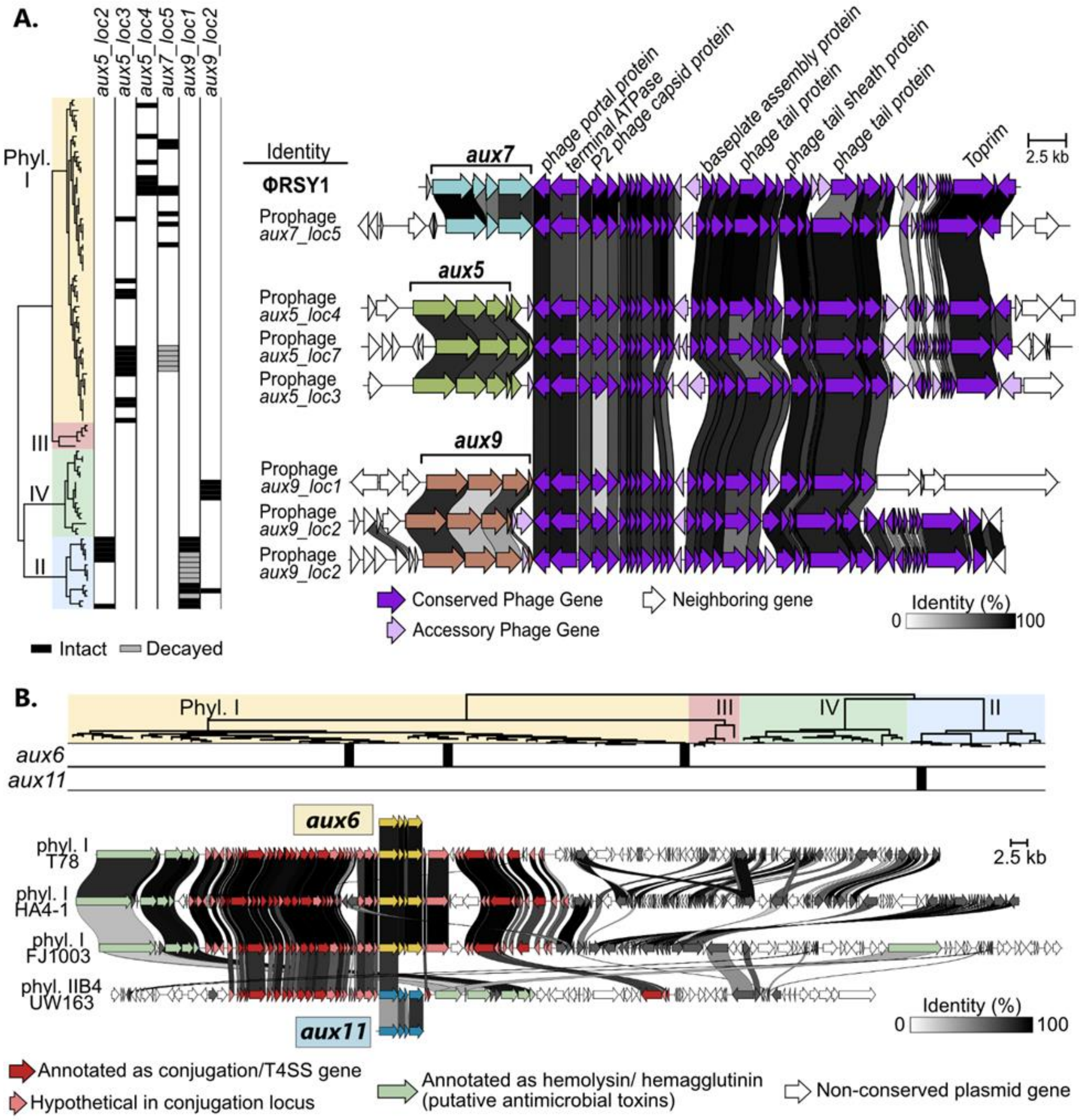


**Figure S31. Mobile genetic elements contribute to horizontal gene transfer of Vgr-linked effector clusters.**

(**A**) *aux5*, *aux7*, and *aux9* are sometimes encoded within RSY1-like bacteriophages. These *aux*-encoding phages insert into 7 different locations and the phylogenetic distribution of each phage-location pair is shown on the left. If the *aux* cluster had clear mutations, the location was marked as decayed (grey). Otherwise, the location is marked as intact (black). Synteny of the phages was visualized with Clinker (right) (**B**) *aux6* and *aux11* are sometimes located on rare, conjugative plasmids. Phylogenetic distribution of *aux*-encoding conjugative plasmids is shown on top. Synteny of the full-length plasmids was visualized with Clinker and genes with putative conjugative functions are highlighted in red.

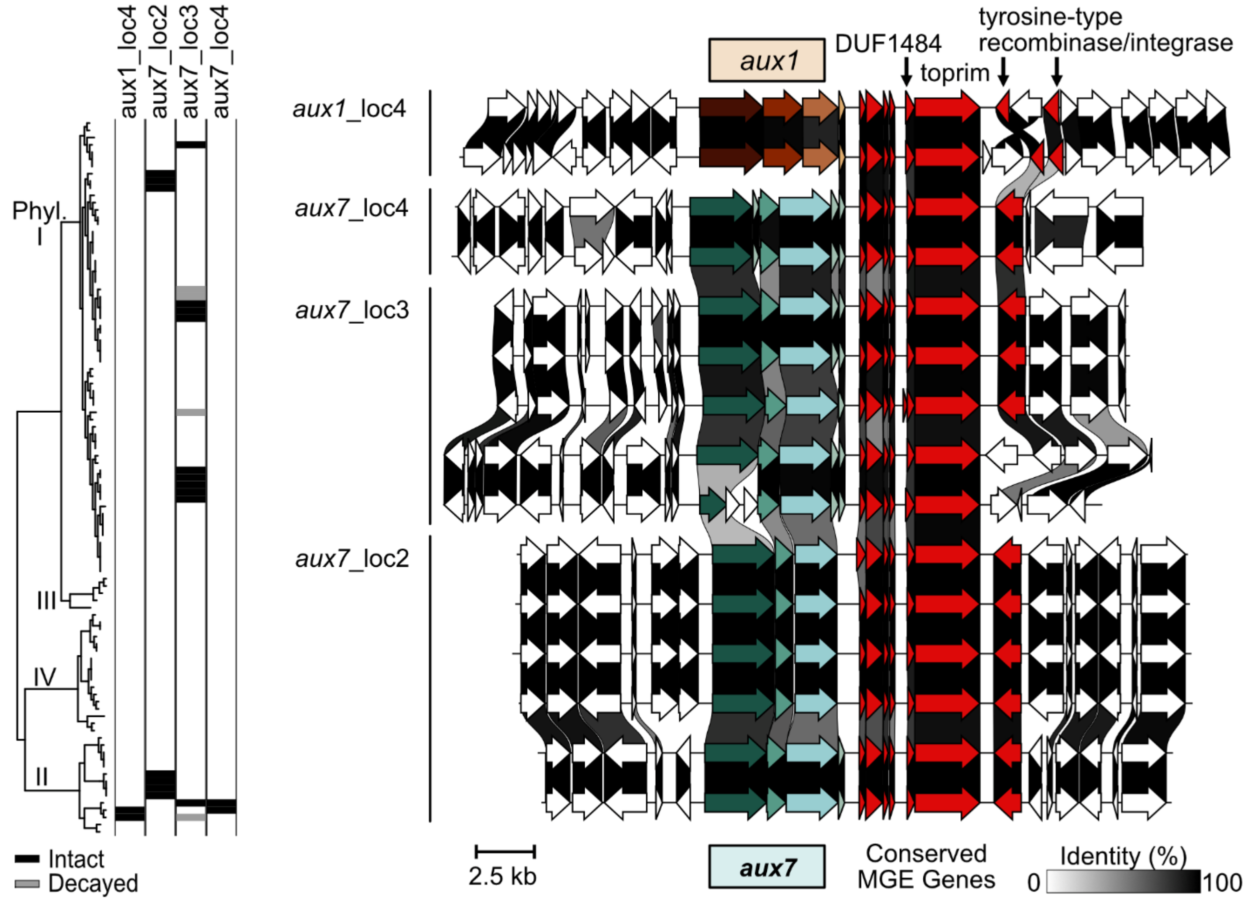


**Figure S32. *aux1* and *aux7* are sometimes located in a toprim/DUF1484 mobile genetic element (MGE).** Synteny was visualized with Clinker. The phylotype-strain name is indicated on the left. *aux11.* Conserved genes annotated with MGE functions are colored red. White genes show the genetic neighborhood where the MGE inserted.

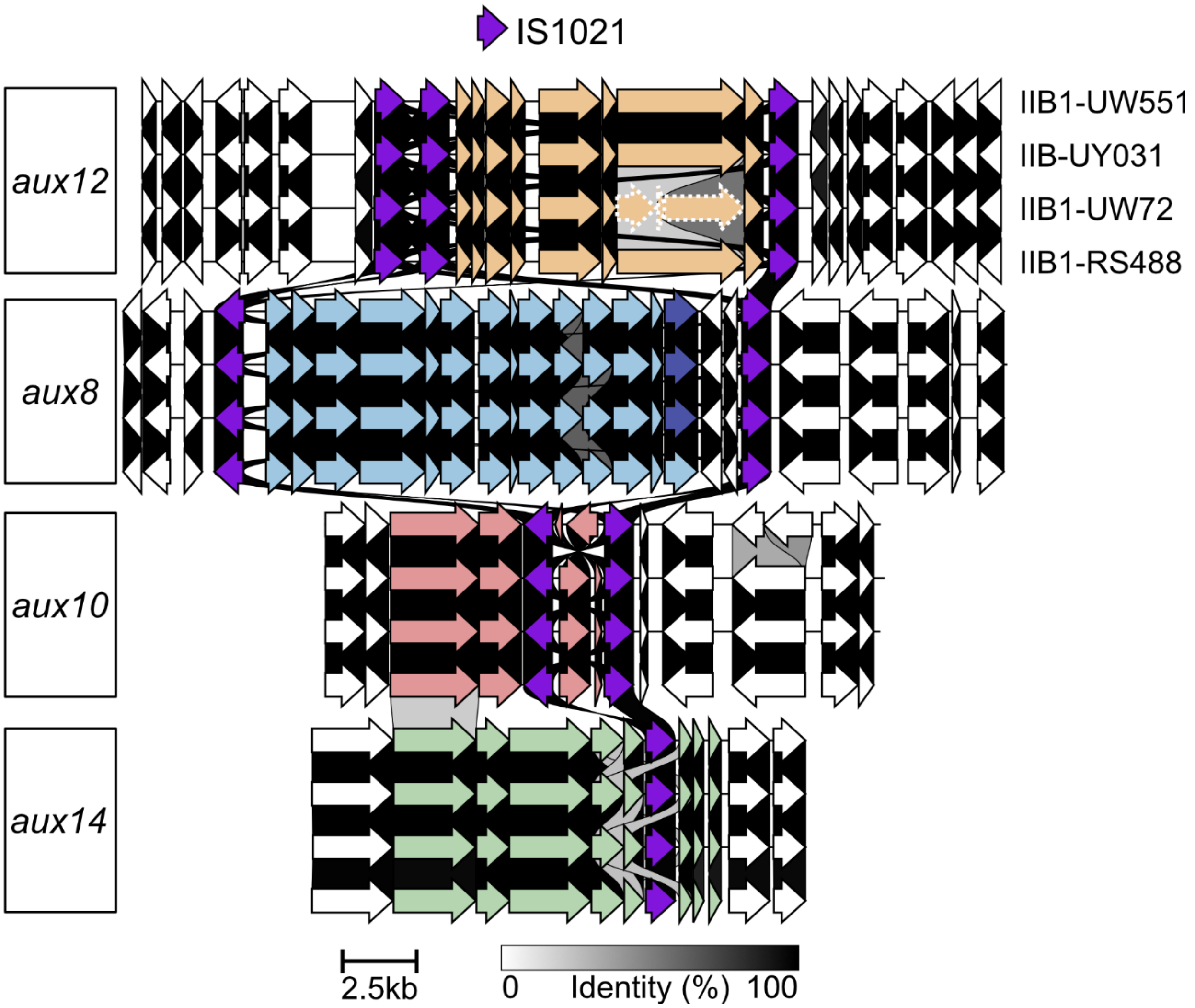


**Figure S33.** ***aux*** **clusters in the pandemic brown rot lineage (IIB1) are associated with IS1021 elements.** Synteny was visualized with Clinker. IS1021 elements are colored red.
